# *De novo* design of modular peptide binding proteins by superhelical matching

**DOI:** 10.1101/2022.11.14.514089

**Authors:** Kejia Wu, Hua Bai, Ya-Ting Chang, Rachel Redler, Kerrie E. McNally, William Sheffler, TJ Brunette, Derrick R. Hicks, Tomos E Morgan, Tim J Stevens, Adam Broerman, Inna Goreshnik, Michelle DeWitt, Cameron M. Chow, Yihang Shen, Lance Stewart, Emmanuel Derivery, Daniel Adriano Silva, Gira Bhabha, Damian Ekiert, David Baker

## Abstract

General approaches for designing sequence-specific peptide binding proteins would have wide utility in proteomics and synthetic biology. Although considerable progress has been made in designing proteins which bind to other proteins, the general peptide binding problem is more challenging as most peptides do not have defined structures in isolation, and to offset the loss in solvation upon binding the protein binding interface has to provide specific hydrogen bonds that complement the majority of the buried peptide’s backbone polar groups (*1*–*3*). Inspired by natural repeat protein-peptide complexes, and engineering efforts to alter their specificity (*4*–*11*), we describe a general approach for *de novo* design of proteins made out of repeating units that bind peptides with repeating sequences such that there is a one to one correspondence between repeat units on the protein and peptide. We develop a rapid docking plus geometric hashing method to identify protein backbones and protein-peptide rigid body arrangements that are compatible with bidentate hydrogen bonds between side chains on the protein and the backbone of the peptide (*12*); the remainder of the protein sequence is then designed using Rosetta to incorporate additional interactions with the peptide and drive folding to the desired structure. We use this approach to design, from scratch, alpha helical repeat proteins that bind six different tripeptide repeat sequences--PLP, LRP, PEW, IYP, PRM and PKW--in near polyproline 2 helical conformations. The proteins are expressed at high levels in E. coli, are hyperstable, and bind peptides with 4-6 copies of the target tripeptide sequences with nanomolar to picomolar affinities both in vitro and in living cells. Crystal structures reveal repeating interactions between protein and peptide interactions as designed, including a ladder of protein sidechain to peptide backbone hydrogen bonds. By redesigning the binding interfaces of individual repeat units, specificity can be achieved for non-repeating sequences, and for naturally occuring proteins containing disordered regions. Our approach provides a general route to designing specific binding proteins for a broad range of repeating and non-repetitive peptide sequences.

## Introduction

A number of naturally occurring protein families bind to peptides with repeating internal sequences (*7, 9*). Particularly well studied are the Armadillo repeat proteins (ARM), such as the nuclear import sequence receptors, which bind to extended peptides with lysine and arginine rich sequences such that each repeat unit in the peptide fits into a repeat unit/module in the protein (*5, 8*). The Plückthun group has demonstrated that the specificity of individual protein repeat units can be re-engineered, enabling broader peptide sequence recognition (*6, 11, 13, 14*). While powerful, this approach is limited to binding peptides in backbone conformations compatible with the geometry of the armadillo repeat. Tetratricopeptide repeat proteins (TPRs) bind peptides with a variety of sequences and conformations, generally with relatively low affinity (∼ µM Kd; for exception see (15)) and with deviations in peptide - protein register which limits their capability of being engineered for more general peptide recognition (*4, 9, 10*).

We set out to generalize peptide recognition by modular repeat-protein scaffolds to arbitrary repeating peptide backbone geometries. This requires solving two main challenges: building protein structures with a repeat spacing and orientation matching that of the target peptide conformation, and ensuring the replacement of peptide-water hydrogen bonds in the unbound state with peptide-protein hydrogen bonds in the bound state. The first challenge is critical for modular and extensible sequence recognition: if individual repeat units in the protein are to bind individual repeat units on the peptide in the same orientation, the geometric phasing of the repeat units on protein and peptide must be compatible. The second challenge is critical for achieving high binding affinity: in conformations other than the alpha and 3-10 helix, the NH and C=O groups make hydrogen bonds with water in the unbound state that need to be replaced with hydrogen bonds to the protein upon binding to avoid incurring a substantial free energy penalty (*16*).

To address the first challenge, we reasoned that a necessary (but not sufficient) criterion for in-phase geometric matching between repeating units on designed protein and peptide was a correspondence between the superhelices that the two trace out. All repeating polymeric structures trace out superhelices which can be described by three parameters: the translation (rise) along the helical axis per repeat unit, the rotation (twist) around this axis, and the distance (radius) of the repeat unit centroid from the axis (Fig. 1A) (*17*). As described in the methods, we generated large sets of repeating protein backbones sampling a wide range of superhelical geometries. We generated corresponding sets of repeating peptide backbones by randomly sampling di-peptide and tri-peptide conformations in allowed regions of the Ramachandran map (avoiding intra-peptide steric clashes), and then repeating these four to six times to generate 12-24 residue peptides. We then searched for matching pairs of repeat protein and repeat peptide backbones, requiring that the rise be within 0.2A, the twist within 5 degrees, and the radius differ by at least 4Å (the difference in radius is necessary to avoid clashing between peptide and protein; the peptide can wrap either outside or inside the protein).

**Figure 1:**
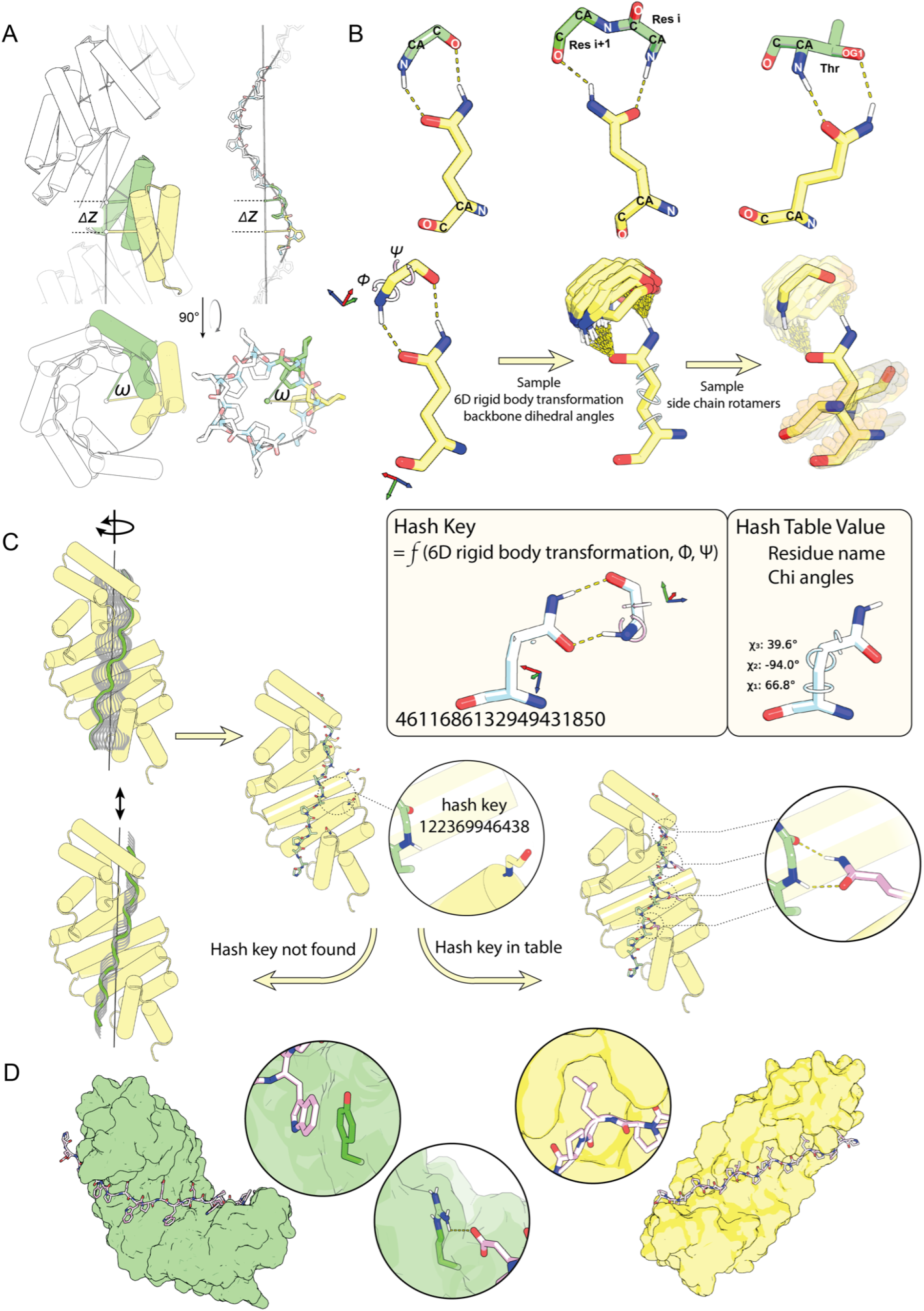
Overview of modular peptide binder design procedure. A) Like all repeating structures, repeat proteins and peptides form superhelices with constant axial displacement (DZ), and angular twist (ω) between adjacent repeat units (shown in green and yellow). For in register binding, the protein and peptide parameters must match (for some integral multiple of repeat units). B) Construction of hash tables for privileged residue-residue interactions. Top row: classes of sidechain backbone interactions for which hash tables were built; (left) sidechain amide group of asparagine or glutamine forming bidentate interactions with the N-H and C=O groups on the backbone of a single residue (left) or consecutive residues (middle), or with the backbone N-H group and sidechain oxygen atom of a serine or threonine (right). Second row: as illustrated for the glutamine - backbone bidentate interaction case, to build the hash table we carry out Monte Carlo sampling over the rigid body orientation between the terminal amide group and the backbone, and the backbone torsions phi and psi, saving configurations with low energy bidentate hydrogen bonds. For each configuration, the possible placements for the backbone of the glutamine are enumerated by growing sidechain rotamers back from the terminal amide. Third row: from the six rigid body degrees of freedom relating the backbones of the two residues, and the phi and psi torsion angles, a hash key is calculated using a 8 dimensional hashing scheme. The hash key is then added to the hash table with the sidechain name and torsions as the value. C) To dock repeat proteins and repeat peptides with compatible superhelical parameters, their superhelical axes are first aligned, and the repeat peptide is then rotated around and slid along this axis. For each of these docks, for each pair of repeat protein-peptide residues within a threshold distance, the hash key is calculated from the rigid body transform between backbones and the backbone torsions of the peptide residue, and the hash table interrogated. If the key is found in the hash table, side chains with the stored identities and torsion angles are installed in the docking interface. D) The sequence of the remainder of the interface is optimized using Rosetta for high affinity binding. Two representative designed binding complexes are shown to highlight the peptide binding groove and the shape complementary. The close-up snapshots illustrate hydrophobic interactions, salt bridges, and π-π stacks incorporated during design.

To address the second challenge, we reasoned that bidentate hydrogen bonds between side chains on the protein and pairs of backbone groups or backbone and sidechain groups on the peptide could allow the burying of sufficient peptide surface area on the protein to achieve high affinity binding without incurring a large desolvation penalty (*18*). As the geometric requirements for such bidentate hydrogen bonds are quite strict, we developed a geometric hashing approach to enable rapid identification of rigid body docks of the peptide on the protein compatible with ladders of bidentate interactions. To generate the hash tables for bidentate sidechain-backbone interactions, Monte Carlo simulations of individual sidechain functional groups making bidentate hydrogen bonding interactions with peptide backbone and/or sidechain groups were carried out using the Rosetta energy function (*12*), and a move set consisting of both rigid body perturbations and changes to the peptide backbone torsions (Fig. 1B; see Methods for details). For each accepted (low energy) arrangement, sidechain rotamer conformations were built backwards from the functional group to identify the possible placements of the protein backbone from which the bidentate interaction could be realized. The results of these calculations were stored in hash tables: for each placement, a hash key was computed from the rigid body transformation and peptide backbone and side chain torsion angles determining the position of the hydrogen bonding groups (for example the phi and psi torsion angles for a bidentate hydrogen bond to the NH and CO groups of the same amino acid), and the chi angles of the corresponding rotamer were stored in the hash for this key (*18*). Hash tables were generated for ASN and GLN making bidentate interactions with the N-H and C=O groups on the backbone of a single residue or adjacent residues, ASP or GLU making bidentate interactions with the N-H groups of two successive amino acids, and for sidechain-sidechain pi-pi and cation-pi interactions (see Methods).

To identify rigid body docks that enable multiple bidentate hydrogen bonds between repeat protein and peptide, we took advantage of the fact that for matching two superhelical structures along their common axis, there are only two degrees of freedom: the translational and rotational offsets of one super helix to the other. For each repeat protein-repeat peptide pair, we carried out a grid search in these two degrees of freedom, sampling relative translations and rotations in ∼1 Å and 10 degree increments (Figure 1E). For each generated dock, we computed the rigid body orientation for each peptide-protein residue pair, and queried the hash tables to very rapidly determine if these were suitable for any of the bidentate interactions; docks for which there were lower than a threshold number of matches were discarded. For the remaining docks, following building of the interacting side chains using the chi angle information stored in the hash, and rigid body minimization to optimize hydrogen bond geometry, we used Rosetta combinatorial optimization to design the protein and peptide sequences (*20*), keeping the residues identified in the hash matching fixed, and enforcing sequence identity between repeats in both peptide and protein (see Methods).

In initial calculations with unrestricted sampling of peptide conformations, designs were generated with a wide range of peptide conformations. Examples of repeat proteins designed to bind to extended beta strand, polypeptide II, and helical peptide backbones, as well as a range of less canonical structures are shown in Fig S1. Reasoning that proline containing peptides would incur a lower entropic cost upon binding, we decided to start experimental characterization with designs containing at least one proline residue; in most such designs the peptide backbone is in or near the polyproline II portion of the Ramachandran map. Our design strategy requires matching the twist of the repeat unit of the peptide with that of the protein, and hence choosing a repeat length of the peptide that generates close to a full 360 degree turn requires less of a twist in the repeat protein; for the polyproline helix there are roughly 3 residues per turn and likely because of this we obtained more designs which target 3 residue than 2 residue proline containing repeat units. We selected for experimental characterization 43 designed complexes with near ideal bidentate hydrogen bonds between protein and peptide, favorable protein-peptide interaction energies (*12*), interface shape complementary (*21*), and few interface unsatisfied hydrogen bonds (*22*) which consistently retained more than 80% of the interchain hydrogen bonds in 20ns molecular dynamics trajectories.

We obtained synthetic genes encoding the designed proteins with a terminal biotinylation tag, expressed the proteins in E. coli, and purified them by Ni-NTA chromatography. 30 of 49 were monomeric and soluble. To assess binding, the target peptides were displayed on the yeast cell surface (*23*), and binding to the repeat proteins was monitored by flow cytometry. To obtain a complete readout of the peptide binding specificity of individual designs, we in parallel used large scale array based oligonucleotide synthesis to generate yeast display libraries encoding all 2 and 3 residue repeat peptides with 8 repeat units each, and used fluorescence activated cell sorting (FACS) followed by Sanger sequencing to identify the peptides recognized by each designed protein. Many of the designs bound peptides with sequences similar to those targeted but the affinity and specificity were both relatively low, with most of the successes for 3 residue repeat units (Fig. S2).

Based on these results, we sought to increase the peptide sequence specificity of the computational design protocol, focusing on design of binders for peptides with 3 residue repeat units. First, we required that each non-proline residue in the peptide make specific contacts with the protein, and that the pockets and grooves engaging sidechains emanating on the two sides of the peptide were quite distinct. Second, following design, we evaluated the change in binding energy (Rosetta DDG) (*24*) for all single residue changes to the peptide repeating unit, and selected only designs for which the design target sequence made the most favorable interactions with the designed protein. Third, we used computational Alanine scanning to remove hydrophobic residues on the protein surface not contributing to binding specificity to decrease non-specific binding (*25*). Fourth, to assess the structural specificity of the designed peptide binding interface, we carried out Monte Carlo flexible backbone docking calculations, starting from large numbers of peptide conformations with superhelical parameters in the range of those of the proteins, and selected those designs with converged peptide backbones (RMSD<2.0 among the top 20 designs with lowest DDG) close to the design model (RMSD<1.5) (Fig. S3).

We tested 54 second-round designed protein-peptide pairs using the yeast flow cytometry assay described above. 42 of the designed proteins were solubly expressed in *E. coli*, and 16 designed bound their targets with considerably higher affinity and specificity than in the first round (Fig. S4). We selected six designs with diverse superhelical parameters and shapes, and a range of target peptides for more detailed characterization (Figure 2). As evident in the design models, these are six repeat proteins (Figure 2A) with a one to one match between repeat units in the protein and in the target peptide (Figure 2B illustrates a single unit interaction). Small Angle X-ray Scattering (SAXS) profiles (*26, 27*) were close to those computed from the design models, suggesting that the proteins fold into the designed shapes in solution (Figure 2C). Circular dichroism (CD) studies showed that all six are largely helical and thermostable up to 95°C, despite that half of the designs don’t fully recover at 20C, which we speculated a small fraction of the proteins likely crash out at 95C due to the exposed surface hydrophobics (Figure 2D). Bio-Layer Interferometry (BLI) characterization of binding to biotinylated target peptides immobilized on Octet sensor chips revealed dissociation constant (Kd) values ranging from <500 pM to ∼40nM; five out of six with dissociation half-life >= 500s (Figure 2E; little decrease in binding was observed after storage of the proteins for 30 days at 4°C, Fig. S5). Three out of the six designs showed little dissociation after 1,000 - 2,000s in buffer, indicating the Kd is too tight to be accurately measured with BLI. The binding surfaces of several related designs were subjected to Site Saturation Mutagenesis (SSM) (*28*) on yeast; and following incorporation of 1-3 enriched substitutions strong binding signals were obtained in flow cytometry using only 10 pM biotinylated cognate peptides for 2 designs (Fig. S6).

**Figure 2:**
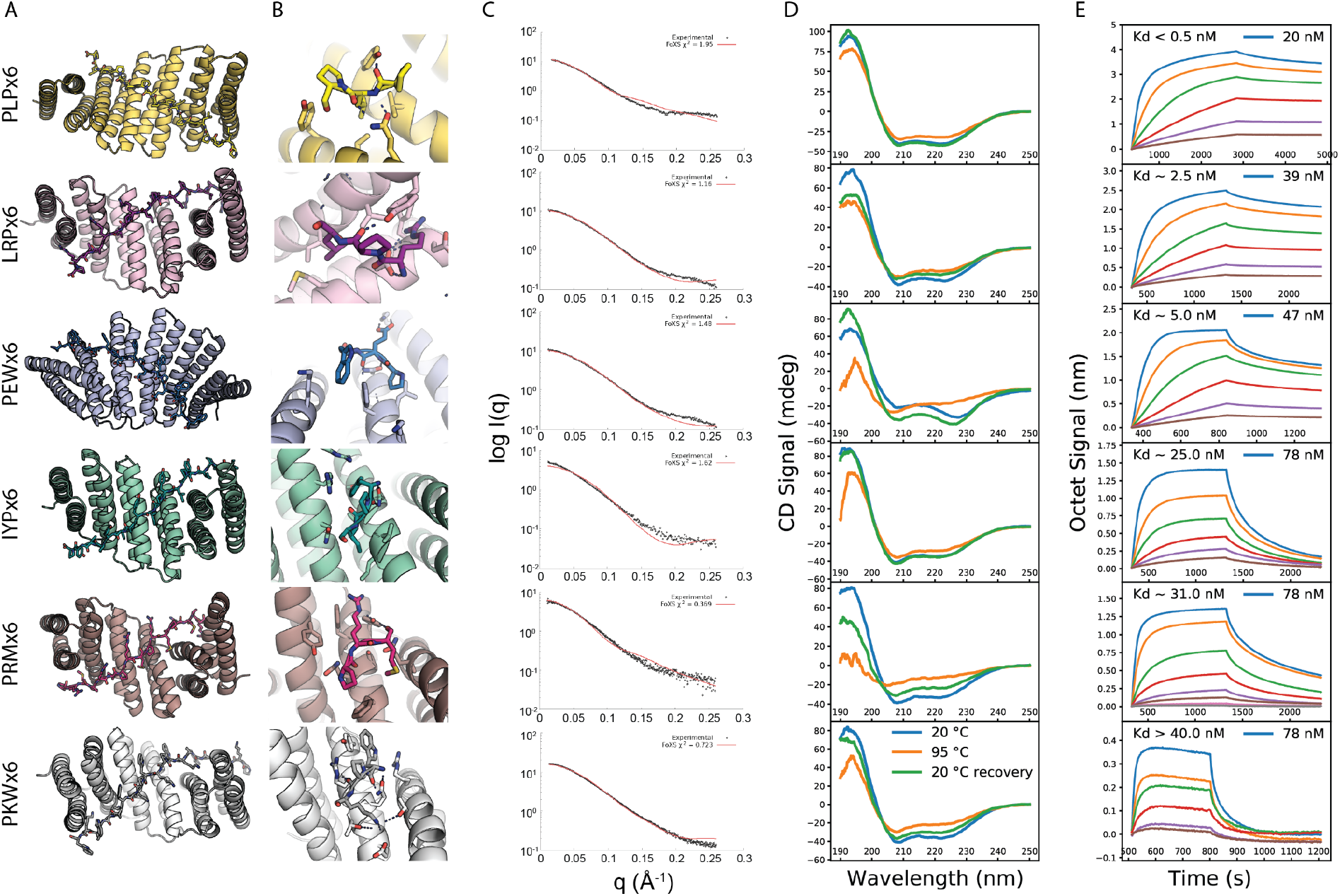
*B*iophysical characterization of designed protein-peptide complexes. **A**. Computational models of the designed six-repeat version of protein-peptide complexes. Designed proteins are shown in cartoons and the peptides in sticks. **B**. Zoom-in view for single designed protein-peptide interaction unit. Residues interacting across the interface are shown in sticks. **C**. predicted SAXS profiles overlaid on experimental SAXS data points. Scattering vector q is on x-axis (from 0 to 0.25), intensity (I) on y axis on log scale. **D**. Circular dichroism spectra at different temperatures (blue: 20 °C, yellow: 95 °C, green: 95 °C followed by 20 °C). **E**. Biolayer interferometry characterization of binding of designed proteins to the corresponding peptide targets. Two-fold serial dilutions were tested for each binder and the highest concentration is labeled. The biotinylated target peptides were loaded onto the Streptavidin (SA) biosensors, and incubated with designed binders in solution to measure association and dissociation.

Many current cell biology approaches (*29*) involve tagging cellular target proteins with a protein or peptide, and then introducing into the same cell a protein which binds the tag with high affinity and specificity, but does not bind endogenous targets. A bottleneck in such studies is that binders obtained from antibody-scaffold (scFV or VHH) based library screens often do not fold properly in the reducing environment of the cytosol, resulting in loss of binding (*30*). We reasoned that our binders would not have this limitation as they are designed for stability and lack disulfide bonds. As a proof of concept, we coexpressed the peptide PLPx6 fused to GFP and its cognate binder, RPB_PLP2_R6, a variant of RPB_PLP1_R6, fused to both mScarlet and a targeting sequence for the mitochondria outer membrane (Fig. 3A). While the PLPx6 peptide on its own was diffuse in the cytosol (Fig. 3B), upon coexpression with the binder, it was relocalized to mitochondria (Fig. 3C; see also Fig. S7 for controls that binder overexpression does not affect mitochondria shape). Thus the PLPx6/RPB_PLP2_R6 pair retains binding activity in cells. Similar results were obtained for IRPx6-GFP and its cognate binder PXX13_FW6 (Fig. 3D,E).

**Figure 3:**
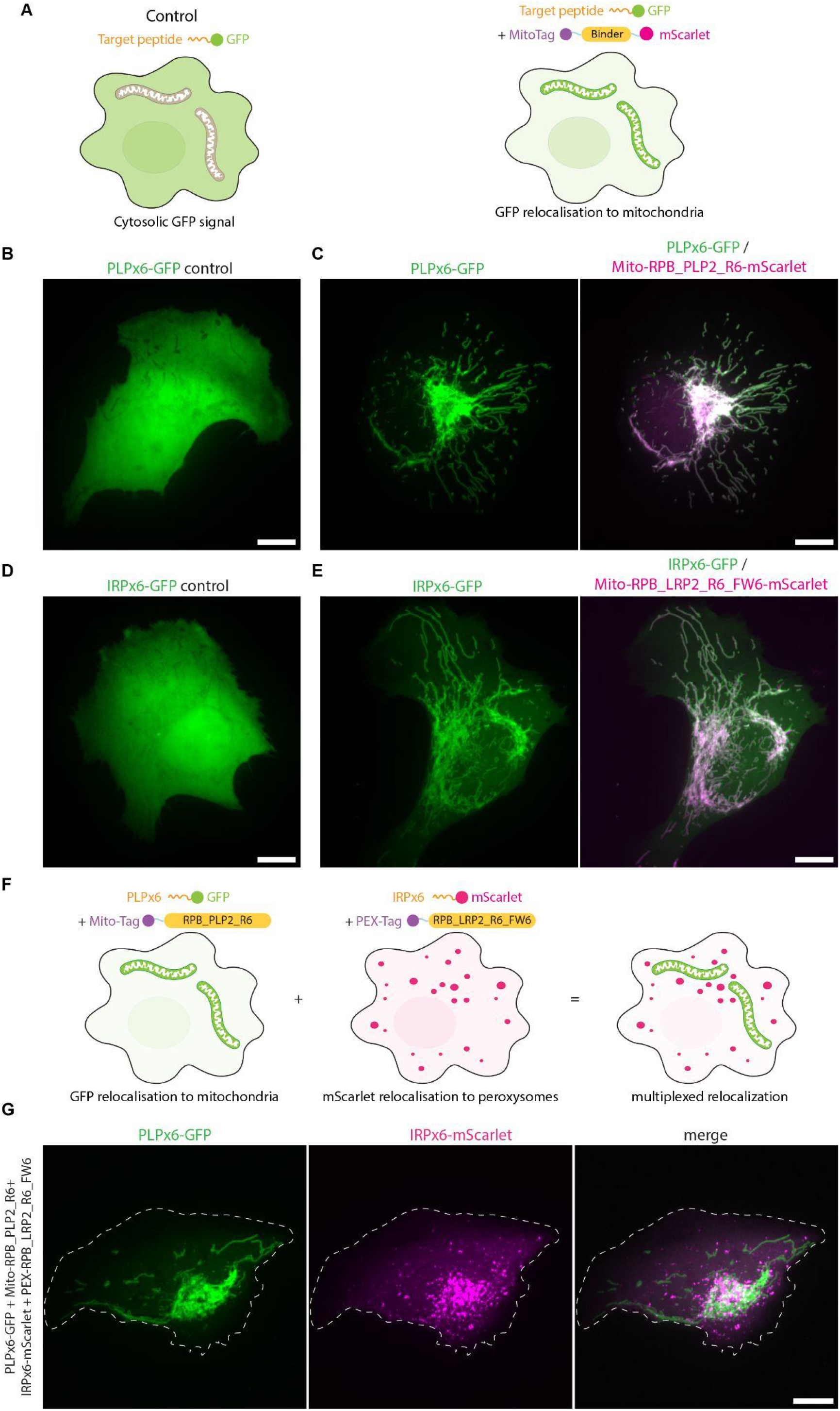
Designed binders function in living cells. **A**. Experimental design: U2OS cells coexpress the target peptide fused to GFP and a fusion between the specific binder fused to mScarlet and a mitochondrial targeting sequence (MitoTag). If binding occurs in cells, the GFP signal is relocalized onto the mitochondria, while control cells not expressing the binder show cytosolic GFP signal. **B-E**. In vivo binding. Live, spreading, U2OS cell expressing PLPx6-GFP alone (**B**), IRPx6-GFP alone (**D**), PLPx6-GFP and Mito-RPB_PLP2_R6-mScarlet (**C**) or IRPx6-GFP and Mito-RPB_LRP2_R6_FW6-mScarlet (**E**) were imaged live by Spinning Disk Confocal Microscopy (SDCM). Note that the GFP signal is cytosolic in control but relocalized to mitochondria upon coexpression with the respective binder. **F-G**. In vivo multiplexing. **F**. Experimental design: cell coexpress two target peptides fused to GFP and mScarlet and their corresponding specific binder fused to mitochondria or peroxysome targetting sequences. If orthogonal binding occurs, GFP and mScarlet signals should not overlap. **G**. Live, spreading, U2OS coexpressing PLPx6-GFP, IRPx6-mScarlet, Mito-RPB_PLP2_R6 and PEX-RPB_LRP2_R6_FW6 imaged by SDCM. Note the absence of overlap between channels. Images correspond to maximum intensity z-projections (Δz= 6 µm). Dash line: cell outline. Scale bars: 10 µm.

If individual repeat units on the designed protein engage individual repeat units on the target peptide, binding affinity should increase with increasing the number of repeats. We investigated this with four of our designed systems, in two cases varying the number of protein repeats while keeping the peptide constant, and in the other two, varying the number of peptide repeats while keeping the protein constant. Six-repeat versions of RPB_LRP2_R6 and RPB_PEW2_R6 had higher affinity for eight-repeat LRP and PEW peptides than four-repeat versions without any decrease in specificity (Fig. S8). Similarly, six-repeat IYP and PLP peptides had higher affinity for six-repeat versions of the cognate designed repeat proteins (RPB_IYP1_R6, RPB_PLP1_R6) than four-repeat versions (Fig. S9). These results are consistent with one to one modular interaction between repeat units on the protein and peptide, and suggest a route to very high binding affinity by simply increasing the number of interacting repeat units. The ability to vary the affinity simply by varying the number of repeats could be useful in many contexts where competitive binding would be advantageous; for example for protein purification by affinity purification, a peptide with a larger number of repeats than that fused to the protein being expressed could be used for elution.

To assess the structural accuracy of our design method, we used X-ray crystallography. We succeeded in obtaining high-resolution co-crystal structures of three first-round designs (RPB_PEW3_R4 - PAWx4, RPB_LRP2_R4 - LRPx4, RPB_PLP3_R6 - PLPx6) and one second-round design (RPB_PLP1_R6 - PLPx6) (Figure 4); and a crystal structure of the unbound first-round design RPB_LRP2_R4 (Fig. S10; interface sidechain RMSDs for all crystal structures are in sup fig x)). In the crystal structure of RPB_PLP3_R6 - PLPx6 design, the PLP units fit exactly into the designed curved groove formed by repeating tyrosine, alanine, and tryptophan residues matching the design model with near atomic accuracy, with Cα rmds of 1.70 Å for the binder apo, 2.00 Å for the peptide neighbor interface and 1.64 Å for the whole complex (Figure 4B, Fig. S11). In the co-crystal structure of RPB_PEW3_R4 - PAWx4, the PAW units bind to a relatively flat groove formed by repeating histidine residues and glutamine residues as designed (Figure 4A, Figure S12); the Cα root-mean-square deviation (RMSD) between design model and crystal structure over the repeat protein is 2.08 Å, and the median value of the RMSD to the crystal structure over the peptide and the binding residues in the flexible docking generated ensemble (which converged less well than for the second round designs) is 2.12Å within 0.03 Å-3.89 Å (Fig. S13). For RPB_LRP2_R4 - LRPx4, flexible backbone docking converged well with the LRP units sitting in between repeating Glutamine residues and Phenylalanine residues as designed, and the peptide Arginine sidechain sampling two distinct states associated with parallel and antiparallel protein binding modes (Fig. S14). The lowest energy docked structure was close to the crystal structure with Cα rmds of 1.15 Å for the binder alone, 0.98 Å for the peptide plus protein contacting residues, and 1.16 Å over the entire complex (Figure 3C, Fig. S14;. SSM binding interface footprinting results were consistent with the design model and crystal structure (Fig. S15), and a FtoW substitution that increases interactions across the interface substantially increases affinity (Fig. S6).

**Figure 4:**
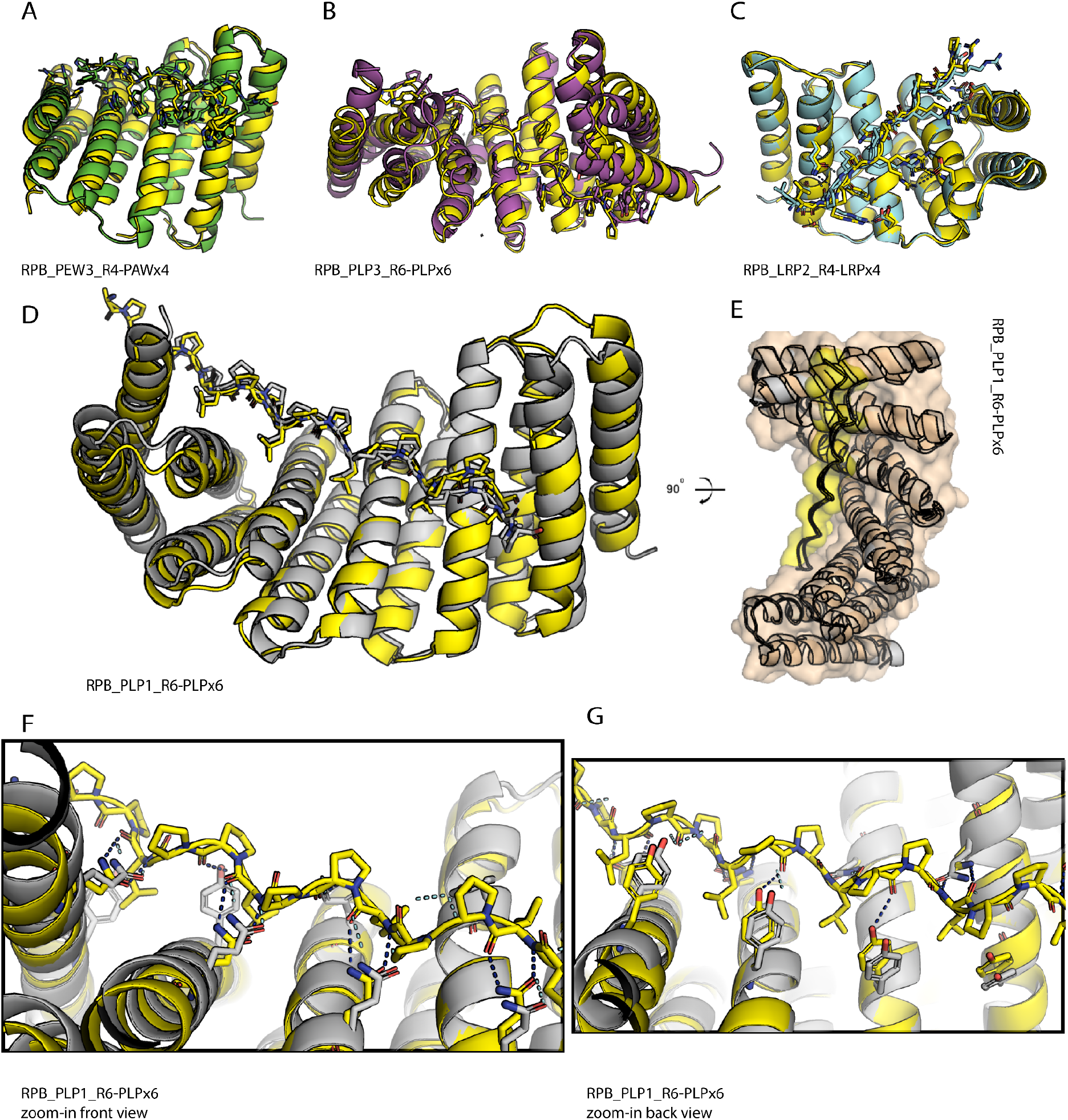
Evaluation of design accuracy by X-ray crystallography. **(A-C)** Superposition of computational design models (colored) on experimentally determined crystal structures (yellow). **A**. RPB_PEW3_R4-PAWx4, **B**. RPB_PLP3_R6-PLPx6, **C**. RPB_LRP2_R4-LRPx4, **D-G**. RPB_PLP1_R6-PLPx6, **D**. overview of superimposition of the computational design model and crystal structure. **E**. 90 degree rotation of **D**.; the complex is shown in surface mode (protein in orange and peptide yellow) for shape complementarity, **F**. Zoom-in interaction of the internal three-unit from **D**. (front view); Glutamine residues from the protein in both design and crystal structure are as sticks to show the accuracy of the designed sidechain-to-backbone bidentate ladder. **G**. Zoom-in interaction of the back view of **F**.; Tyrosine residues from the protein in both design and crystal structure are in sticks to show the accuracy of designed polar interactions on the other side.

The 2.15A crystal structure of the 2nd round design RPB_PLP1_R6 - PLPx6 highlights key features of the computational design protocol. The PLPx6 peptide binds to the slightly curved groove primarily through polar interactions from tyrosine, hydrophobic interactions from Valine, and sidechain-backbone bidentate hydrogen bonds from Glutamine exactly as designed (Figure 4D-4G). The Cα rmds are 1.11 Å for the peptide neighbor interface and 1.81 Å for the binder apo, 1.91 Å for the complex. All interacting side-chains from both the protein side and the peptide side in the computational design model are nearly perfectly recapitulated in the crystal structure. This design has near picomolar binding affinity (Figure 2D) and high specificity for the PLP target sequence (Figure 5A).

**Figure 5:**
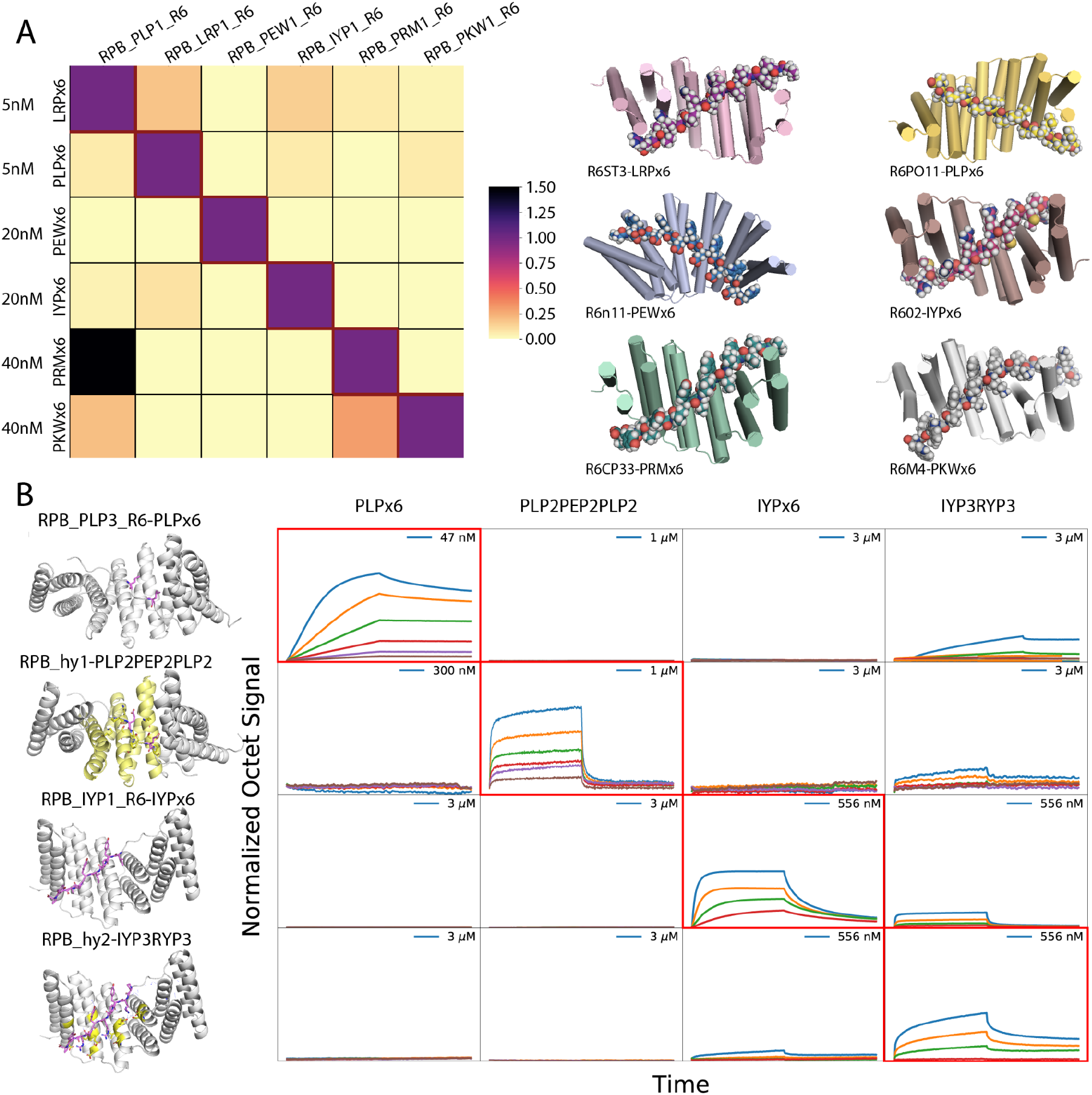
Designed protein-peptide interaction specificity. **A**. (left) To assess the cross reactivity of each designed peptide binder in Fig 2 with each target peptide, biotinylated target peptides were loaded onto biolayer interferometry SA sensors, allowed to equilibrate, and baseline signal set to zero. The BLI tips were then placed into solution containing proteins at the indicated concentrations for 500 seconds, washed with buffer, and dissociation was monitored for an additional 500 seconds. The heatmap shows the maximum signal for each binder-target pair normalized by the maximum signal of the cognate designed binder-target pair. (Right) Surface shape complementarity of the cognate complexes; peptides are in sphere representation. **B**. Modular sequence design generates binders for not strictly repeating peptide sequences. (left, row 1 and 2) Binding of base complex RPB_PLP3_R6-PLPx6 and hybrid binder complex RPB_hy1-PLP_2_PEP_2_PLP_2_. (left, row 3 and 4) Binding of base complex RPB_IYP1_R6-IYPx6 and hybrid binder complex RPB_hy2-IYP_3_RYP_3_. The redesigned peptide and protein residues are shown in purple sticks and yellow respectively. (right) orthogonality matrix: Biotinylated target peptides were loaded onto biosensors, and incubated with designed binders in solution at the indicated concentrations. Red rectangle box indicates cognate complexes. Octet signal was normalized by the maximum signal of the cognate designed binder-target pair.

We next investigated the specificity of the six designs (Fig 5A). The PLPx6, LRPx6, PEWx6, IYPx6, PKWx6 binders showed almost complete orthogonality in the 5∼40nM concentration range, with each design binding its cognate designed repeat peptide much more strongly than the other repeat peptides. For example, PLPx6 binds design RPB_PLP1_R6 strongly at 5nM, but shows no binding signal to design RPB_IYP1_R6 at 40nM, while PEWx6 binds design RPB_PEW1_R6 but not design RPB_PKW1_R6 at all at 20nM. Some crosstalk was observed between the PRMx6 and LRPx6 binders perhaps involving the arginine residue which makes cation-pi interactions in both designs. We observe similar orthogonality of the interaction between peptide/binder pairs in cells, as the IRPx6 and PLPx6 binders specifically direct localization of their cognate peptides to different compartments when coexpressed in the same cells (Fig. 3E,F). By enabling the design of multiple orthogonal protein-peptide pairs, our approach provides a route to probing the effects of localizing different proteins to different locations in the same cell.

As described thus far, our approach enables specific binding of peptides with perfectly repeating structures. To go beyond this limitation and enable targeting of a much wider range of non-repeating peptides, we investigated the redesign of a subset of the peptide repeat unit binding pockets to change their specificity. We broke the symmetry in the designed repetitive binding interface by redesigning both protein and peptide in one or more repeats of six-repeat complexes; the rest of the interface was kept untouched to maintain binding affinity. Following redesign, the peptide backbone conformation was optimized by Monte Carlo resampling and rigid body optimization (see Methods). Designs were selected for experimental characterization as described above, favoring those for which the new design had lower binding energy for the new peptide than the original peptide.

We redesigned the PLPx6 binder RPB_PLP3_R6 to bind two PEP units in the third and fourth positions (target binding sequence PLPPLPPEPPEPPLPPLP, or more concisely, PLP_2_PEP_2_PLP_2_). The redesigned protein, called RPB_hyb1_R6, bound the redesigned peptide considerably more tightly in octet experiments, while the original design favored the original perfectly repeating sequence, resulting in nearly complete orthogonality (Figure 5B). We next designed another hybrid starting from the RPB_IYP1_R6 - IYPx6 complex, changing 3 of the IYP units to RYP, generating IYP_3_RYP_3_, and redesigning the corresponding binding pockets. The new design, RPB_hyb2_R6, selectively bound the intended cognate target as well (Figure 5C). We measured binding of all four proteins against all four peptides, and observed quite high specificity of the designed repeat proteins for their intended peptide targets (Figure 5B-5C).

The ability to design hybrid binders against non-repetitive sequences opens the door to the *de novo* design of binders against endogenous proteins. Intrinsically disordered regions (IDR) are targets of choice, as they have been very difficult to specifically recognize using other approaches, and folding will not interfere with binding. As a proof of concept, we focused on human ZFC3H1, a 200 kDa protein that together with MTR4 forms the heterotetrameric poly(A) tail exosome targeting (PAXT) complex, which directs a subset of long polyadenylated poly(A) RNAs for exosomal degradation (Fig. 6A and ref(*31, 32*)). We designed binders against ZFC3H residues 594-620 (PLP_4_PEDPEQPPKPPF) which lie within a ∼100 residue disordered region (Fig. 6A), by extending both the protein and peptide in the PLPx4 designed complex. On the peptide side, we kept the (PLP)x4 backbone fixed, and used Monte Carlo sampling with Ramachandran map biases to model the remaining sequence (PEDPEQPPKPPF); on the protein side, we extended the PLPx4 design with four additional repeats and designed the interface with each peptide conformer, and selected eight designs for experimental characterization, as described above for the pure repeat binders. Eight designs were expressed, and seven found to bind the extended target peptide by biolayer interferometry (Fig. S19). The two highest affinity designs were further characterized by fluorescence polarization and found to bind the 24 residue target peptide (Fig. 5C). The one with the highest affinity, named αZFC-high (Fig. 6B), also co-eluted with a 103 amino acid segment of the disordered region of ZFC3H1 containing the targeting sequence by Size-Exclusion Chromatography (SEC) (Fig. 6D), demonstrating that the binder can recognize the target peptide in a larger protein context. αZFC-high specifically pulled down the endogenous ZFC3H1 from human cell extracts when assessed by western blot with established antibodies (Fig. 6E, upper panel), in contrast to the lower-affinity binder αZFC-low, which has similar size and surface composition and hence provides a control for non-specific association (see Fig. S16 for replicates, and Fig.6F for independent identification of ZFC3H1 by mass spectrometry). Mass Spectrometry revealed that MTR4 was enriched in the αZFC-high pull down, demonstrating that the binder can recognise the native PAXT complex in a physiological context. We also detected in the αZFC-high pulldown, but not the αZFC-low pulldown, additional ZFC3H1 partners present in the Bioplex 3.0 interactome in multiple cell lines (*33, 34*), including BUB3 and ZN207, and multiple RNA binding proteins which likely associate with PAXT - RNA assemblies (Fig.6F see supplementary table 5 for full proteomics dataset).

**Figure 6:**
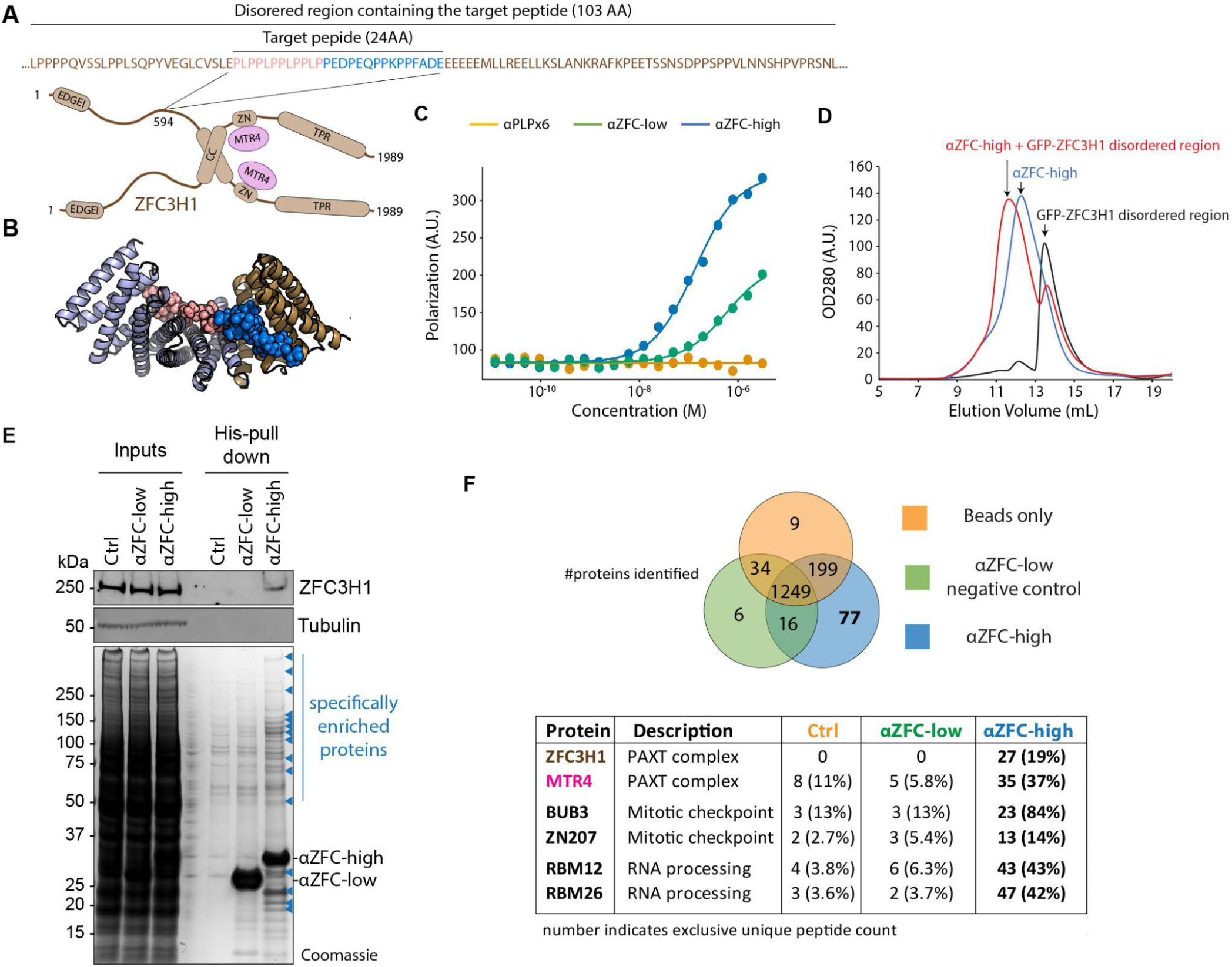
Design of binders specific for endogenous human proteins. **A**.Schematic model of the human PAXT complex composed of a heterotetramer of ZFC3H1 and MTR4. Domain acronyms: CC: Coiled-coil ; ZN: Zn-finger domain. Inset shows the details of the environment of the target sequence. **B**. Surface shape complementarity between the target peptide from ZFC3H1 (sphere) and the highest affinity cognate binder αZFC-high. **C**. Fluorescence polarization binding curves between indicated ZFC3H1 binders and the target ZFC3H1 peptide (PLP)4PEDPEQPPKPP. As a negative control, we used the (PLP)x6 binder, RPB_PLP3_R6 (see Fig.4). αZFC-high shows higher binding affinity to the target peptide than αZFC-low, on the contrary to RPB_PLP3_R6, which shows negligible binding. **D**. Superdex 200 10/300 GL size exclusion chromatography profiles of purified αZFC-high, a fusion between GFP and a 103AA fragment of the disordered region of ZFC3H1 containing the target sequence (see A.), or a 1:1 mix of the two after 2h incubation. **E**. Top: Hela cell extracts were subjected to pull down using indicated binders bound to NiNTA agarose beads, or naked beads as control. Recovered proteins were processed for western blot against endogenous ZFC3H1 (or tubulin as a loading control). Bottom, coomassie stained, SDS-PAGE gel of the samples analyzed in top panel. These panels are representative of n=3 experiments. **F**. Proteomics analysis of the his-pull down samples shown in C. Top panel: overlap between the proteins identified, setting s threshold of five peptides for correct identification. bottom panel: examples of proteins identified (number indicates exclusive peptide counts. protein coverage is indicated in parenthesis). See Supplementary Table 5 for the full dataset.

## Conclusion

Our results demonstrate that by matching superhelical parameters between repeating protein and peptide conformations together with incorporation of specific hydrogen bonding and hydrophobic interactions, new repeat proteins binding repeating peptide sequences with high affinity and specificity can now be designed. The approach should be generalizable to a wide range of repeating peptide structures, and the ability to break symmetry by redesigning individual repeat units opens the door to more general peptide recognition. Our approach complements current efforts at achieving general peptide recognition by redesign of naturally occurring repeat proteins; an advantage of our approach is that a much broader range of protein conformations and binding site geometries can be generated by de novo protein design than by starting with a native protein backbone. Proteins binding repeating or nearly repeating sequences could have applications as affinity reagents for diseases such as Huntington’s which are associated with repeat expansions, and rigid fusion of protein modules designed, using the approach described here, to recognize different di, tri and tetra peptide sequences provides an avenue to achieving specific recognition of entirely non-repeating sequences. The ability to design specific binders to proteins containing large disordered regions, demonstrated by the specific pull down of the PAXT complex (Fig 6), should contribute to delineating the functions of this important but relatively poorly understood class of proteins and reduce reliance on animal immunization to generate antibodies, which can also suffer from reproducibility issues. More generally, our results demonstrate the power of computational protein design for targeting peptides not having rigid three dimensional structures, and as the designed proteins are expressed at quite high levels and very stable, we anticipate that these and further designs for a wider range of target sequences should find broad use in proteomics and other applications requiring specific peptide recognition.

## Funding

This work was supported by The Audacious Project at the Institute for Protein Design (D.B., K.W., M.D., D.A.S., A.B.), The Michelson Found Animals Foundation Grant Number GM15-S01 (L.S., K.W., D.B.), the National Institute on Aging grant 5U19AG065156-02 (D.H., K.W., D.B.), the Howard Hughes Medical Institute (D.B., W.S., H.B.), The Open Philanthropy Project Improving Protein Design Fund (A.C., R.R., C.M.C., G.B., D.E., D.B.), The Donald and Jo Anne Petersen Endowment for Accelerating Advancements in Alzheimer’s Disease Research (T.J.B., D.B.), a donation from AMGEN to the Institute for Protein Design (I.G.), the Medical Research Council (MC_UP_1201/13 to E.D., T.E.M, T.J.S.), the Human Frontier Science Program (CDA00034/2017-C to E.D.) and a Sir Henry Wellcome Postdoctoral Fellowship (K.M.).

## Competing Interests

Each contributor attests that they have no competing interests relating to the subject contribution, except as disclosed. K.W., H.B., D.R.H., T.J.B., K.E.M., T.J.S, T.E.M, A.C., R.R., G.B., D.E., L.S., E.D, D.A.S., W.S., I.G. and D.B. are co-inventors on a patent application that incorporate discoveries described in this article.

We thank B. Wicky, A. Ljubetic, and I. Lutz for advice on the split luciferase assay for the second-round design screening, C. Xu for help trouble-shooting experiments, T. Schlichtharle for discussion, L. Cao and I. Goreshnik for advice on biolayer interferometry, H. Pyles for advice on circular dichroism (CD) and designed helical repeat proteins (DHR),Ramanujan Hegde for the suggestion to target disordered regions of endogenous proteins, K. Van Wormer and A. Curtis Smith for laboratory support during COVID-19.

## Author contributions

K.W. and H.B. contributed equally to this work; K.W., D.A.S. and D.B. designed the research; D.A.S. and D.B. developed the preliminary computational method and hash database; W.S. contributed to the hash database development; K.W. updated the computational method with the help from D.A.S and H.B.; H.B. updated the hash database to be more general; Y.S. helped and contributed to the first hash database development; K.W. and T.J.B. designed the polyproline 2 DHR scaffold library using the method developed by D.R.H.; K.W. designed the binders with the help from H.B; H.B. and K.W. performed the yeast screening, expression and binding experiments with the help from I.G. for the first-round design characterization; K.W. performed biolayer interferometry, Octet assays for the second-round design characterization; H.B. constructed and screened site saturation mutagenesis libraries (SSMs). A.C., R.R., G.B., D.E. solved the structures of RPB_PEW3_R4-PEWx4, RPB_PLP3_R6-PLPx6, RPB_LRP2_R4-LRPx4 and RPB_PLP1_R6-PLPx6; K.E.M. designed and performed all cell experiments in this work, in particular the multiplex binding assay and the demonstration of the endogenous binder for ZFC3H1. E.D. identified ZFC3H1 as a good target for the development of an endogenous binder with help from T.J.S. T.E.M. performed mass spectrometry analysis ; A.B. helped with the modular binding assay; M.D. and C.M.C. helped with preparing protein samples for crystallography. All authors analyzed data. L.S., D.A.S. and D.B. supervised research. K.W. and D.B. wrote the manuscript with the input from the other authors. All authors revised the manuscript.

