## Supplemental Figures for "*De novo* design of modular peptide binding proteins by superhelical matching"

**This PDF file includes:**

Materials and Methods
Figs. S1 to S15
Tables S1 to S3
Caption for Table S4

**Materials and Methods**

Synthetic gene constructs

All genes in this work were ordered from either Integrated DNA Technologies (IDT) or Genscript. For both the first and second round designs, a His-tag containing TEV protease cleavage site and short linkers were added to the N-terminus of protein sequences. For the protein lacking a Tryptophan residue, a single Tryptophan was added to the short N-terminal linker following the TEV protease cleavage site to help with protein concentration quantification by A280. The protein sequence along with linker (MGSSHHHHHHHHSSGGSGGLNDIFEAQKIEWHEGGSGGSENLYFQSG or LEHHHHHH) was reverse translated into DNA using a custom python script that attempts to maximize host-specific codon adaptation index[1] and IDT synthesize-ability, which includes optimizing whole gene and local GC content as well as removing repetitive sequences. Finally, a TAATCA stop codon was appended to the end of each gene. Genes were delivered cloned into pET-29b+ between NdeI/XhoI restriction sites. For the second round designs, the designed amino acid sequences were inserted into pET-29b+ between NdeI/XhoI restriction sites directly.

For the ZFC3H1-103 disordered region, the 103 amino acids harboring the key targeting sequence (LPPPPQVSSLPPLSQPYVEGLCVSLEPLPPLPPLPPLPPEDPEQPPKPPFADEEEEEE MLLREELLKSLANKRAFKPEETSSNSDPPSPPVLNNSHPVPRSNL) was cloned into a customized vector with sfGFP at N-terminal and His6 at C-terminal with linker (GGSGSG) in between.

Protein expression and purification

Proteins were transformed into Lemo21(DE3) E. coli from New England Biolabs (NEB) and then expressed as 50 ml cultures in 250 ml flasks using Studiers M2 autoinduction media with 50 ug/mL kanamycin. The cultures were either grown at 37°C for ~6-8 hours and then ~18°C overnight (~14 hours) or at 37°C the entire time ~14 hours. Cells were pelleted at 4,000g for 10 minutes, after which the supernatant was discarded. Pellets were resuspended in 30 ml lysis buffer (25 mM Tris HCl pH 8, 150 mM NaCl, 30 mM imidazole, 1mM PMSF, 0.75% CHAPS, 1 mM DNase, 10mM Lysozyme, with Thermo Scientific Pierce protease inhibitor tablet). Cell suspensions were lysed by microfluidizer or sonication, and the lysate was clarified at 20,000g for ~30 minutes. The His-tagged proteins were bound to Ni-NTA resin (Qiagen) during gravity flow and washed with a wash buffer (25 mM Tris HCl pH 8, 150 mM NaCl, 30 mM imidazole). Protein was eluted with an elution buffer (25 mM Tris HCl pH 8, 150 mM NaCl, 300 mM imidazole). For the first round designs, the His-tag was removed by TEV cleavage, followed by IMAC purification to remove TEV protease. The flowthrough was collected and concentrated prior to further purification by SEC/FPLC on a superdex 200 increase 10/300 GL column in TBS (25 mM Tris pH 8.0, 150 mM NaCl).

Circular dichroism

Circular dichroism spectra were measured with an AVIV Model 420 DC or Jasco J-1500 CD spectrometer. Samples were 0.25 mg/mL in TBS (25 mM Tris pH 8.0, 150 mM NaCl), and a 1-mm path length cuvette was used. The CD signal was converted to mean residue ellipticity by dividing the raw spectra by N × C × L × 10, where N is the number of residues, C is the concentration of protein, and L is the path length (0.1 cm).

Size exclusion chromatography with multi-angle light scattering

Purified samples after the initial SEC run, samples were pooled then concentrated or diluted as needed to a final concentration of 2 mg/mL. 100 uL of each sample was then run through a high-performance liquid chromatography system (Agilent) using a Superdex 200 10/300 GL column. These fractionation runs were coupled to a multi-angle light scattering detector (Wyatt) in order to determine the absolute molecular weights for each designed protein as described previously[2].

Small angle X-ray scattering

Small-Angle X-ray Scattering (SAXS) was collected at the SIBYLS High Throughput SAXS Advanced Light Source in Berkeley, California[3]. Beam exposures of 0.3 s for 10.2 s resulted in 33 frames per sample. Data was collected at low (~1.5 mg/mL) and high (~2-3 mg/mL) protein concentrations in SAXS buffer (25mM Tris pH 8.0, 150mM NaCl, 2% glycerol). The sibyls website(“SAXS FrameSlice” n.d.) was used to analyze the data for high and low centration samples and average the best dataset. If there was obvious aggregation over the 33 frames, only the data points before aggregation arose were used in the Gunier region, otherwise, all data was included for the Gunier region. All data was used for Porod and Wide regions. The averaged file was used with scatter.jar to remove data points with outlier residuals in the Gunier region. Finally, the data was truncated at 0.25 q. This dataset was then compared to the predicted SAXS profile based on the design model using the FoxS SAXS server(“FoXS Server: Fast X-Ray Scattering” n.d.), and volatility ratio (Vr) was calculated to quantify how well the predicted and data matched the experimental data. Proteins with Vr of less than 2.5 were considered to be folded to the designed quaternary shape.

Biolayer interferometry

Biolayer interferometry binding data were collected in an Octet RED96 (ForteBio) and processed using the instrument’s integrated software. To measure the affinity of peptide binders, N-terminal biotinylated (biotin-Ahx) target peptides with a short linker (GGS) were loaded onto streptavidin-coated biosensors (SA ForteBio) at 50-100 nM in binding buffer (10 mM HEPES (pH 7.4), 150 mM NaCl, 3 mM EDTA, 0.05% surfactant P20, 0.5% non-fat dry milk) for 120 s. Analyte proteins were diluted from concentrated stocks into the binding buffer. After baseline measurement in the binding buffer alone, the binding kinetics were monitored by dipping the biosensors in wells containing the target protein at the indicated concentration (association step) and then dipping the sensors back into baseline/buffer (dissociation).

Yeast surface display

S. cerevisiae EBY100 strain cultures were grown in C-Trp-Ura media and induced in SGCAA media following the protocol in (reference). Cells were washed with PBSF (PBS with 1% BSA) and labeled with biotinylated designed proteins using two labeling methods, with-avidity and without-avidity labeling. For the with-avidity method, the cells were incubated with biotinylated RBD, together with anti-c-Myc fluorescein isothiocyanate (FITC, Miltenyi Biotech) and streptavidin– phycoerythrin (SAPE, ThermoFisher). The concentration of SAPE in the with-avidity method was used at ¼ concentration of the biotinylated RBD. The with-avidity method was used in the first few rounds of screening against the repeat peptide library to fish out weak binder candidates. For the without-avidity method, the cells were firstly incubated with biotinylated designed proteins, washed, secondarily labeled with SAPE and FITC.

Crystallography

**Crystallographic Data Collection and Refinement Statistics**

| | RPB_PEW3_R4-PAWx4 | RPB_PLP3_R6-PLPx6 | RPB_LRP2_R4-LRPx4 | RPB_PLP1_R6-PLPx6 |
|---|---|---|---|---|
| Data collection | | | | |
| Beamline | APS 23-ID-D | APS 23-ID-D | APS 23-ID-B | APS 23-ID-B |
| Wavelength | 1.033 | 1.033 | 1.033 | 1.033 |
| Space group | $P2_12_12_1$ | *I422* | $P3_221$ | *R32* |
| Cell dimensions | | | | |
| *a*, *b*, *c* (Å) | 45.4, 67.6, 71.3 | 124.5, 124.5, 105.0 | 70.8, 70.8, 128.2 | 99.4, 99.4, 173.2 |
| *α*, *β*, *γ* (°) | 90.0, 90.0, 90.0 | 90.0, 90.0, 90.0 | 90.0, 90.0, 120.0 | 90.0, 90.0, 120.0 |
| Resolution (Å) | 38.3 - 2.70 (2.80-2.70)[1] | 45.1-2.68 (2.78-2.68)[1] | 35.4-3.18 (3.92-3.18)[1] | 37.7-2.15 (2.21-2.15)[1] |
| Total Observations | 39,005 | 155,310 | 64,884 | 186,166 |
| Unique reflections | 6,383 | 11,891 | 6,621 | 18,220 |
| Redundancy | 6.1 (6.4)[1] | 13.1 (13.3)[1] | 9.8 (10.2)[1] | 10.2 (10.5)[1] |

| | | | | |
|---|---|---|---|---|
| Completeness (%) | 99.6 (99.8)[1] | 92.1 (77.4)[1] | 89.1 (67.9)[1] | 99.9 (100)[1] |
| $R_{meas}$ | 0.08 (1.14)[1] | 0.11 (2.51)[1] | 0.08 (3.38)[1] | 0.06 (2.45)[1] |
| $I/\sigma(I)$ | 12.0 (1.27)[1] | 18.5 (1.21)[1] | 14.1 (0.69)[1] | 18.1 (0.85)[1] |
| $CC_{1/2}$ | 1.00 (0.81)[1] | 1.00 (0.51)[1] | 1.00 (0.69)[1] | 1.00 (0.45)[1] |
| Refinement | | | | |
| Resolution (Å) | 38.3 - 2.70 | 45.1 - 2.68 | 35.4 - 3.18 | 37.7 - 2.15 |
| Reflections (work) | 6,057 | 10,412 | 5,357 | 15,249 |
| Reflections (test) | 316 | 547 | 294 | 1,672 |
| $R_{work}$ / $R_{free}$ (%) | 24.5 / 27.0 | 22.6 / 27.7 | 20.7 / 25.6 | 23.2 / 27.6 |
| Average B-factor ($Å^2$) | 114.9 | 86.9 | 162.9 | 86.9 |
| No. atoms | | | | |
| Protein | 1,502 | 2,375 | 1,372 | 2,172 |
| Peptide | 104 | 139 | 104 | 240 |
| Water | 0 | 0 | 0 | 7 |
| Ramachandran[2] | | | | |
| Favored (%) | 98.0 | 98.4 | 97.2 | 99.3 |
| Outliers (%) | 0.0 | 0.0 | 0.6 | 0.0 |
| Rotamers[2] | | | | |
| Favored (%) | 96.20 | 98.8 | 94.08 | 97.1 |
| Outliers (%) | 0.0 | 0.0 | 0.0 | 0.0 |
| R.m.s. deviations | | | | |
| Bond lengths (Å) | 0.002 | 0.002 | 0.004 | 0.002 |
| Bond angles (°) | 0.41 | 0.49 | 0.62 | 0.41 |
| Molprobity[2] | | | | |
| Molprobity Score | 1.10 | 1.38 | 1.78 | 1.01 |
| Percentile | 100th | 100th | 100th | 100th |
| Clashscore | 3.12 | 6.99 | 13.35 | 3.06 |
| Percentile | 100th | 99th | 95th | 99th |
| PDB ID | 7UDJ | 7UE2 | 7UDK | 7UDL |

| | RPB_PLP1_R6-PLPx6, alternative conformation 1 | RPB_PLP1_R6-PLPx6, alternative conformation 2 | RPB_LRP2_R4, pseudopolymeric |
|---|---|---|---|
| Data collection | | | |
| Beamline | APS 23-ID-B | APS 23-ID-B | APS 23-ID-B |
| Wavelength | 1.033 | 1.033 | 1.033 |
| Space group | $P22_12_1$ | $P22_12_1$ | $P3_221$ |
| Cell dimensions | | | |
| *a*, *b*, *c* (Å) | 54.4, 80.3, 154.4 | 80.4, 86.5, 110.2 | 72.6, 72.6, 127.3 |
| *α*, *β*, *γ* (°) | 90.0, 90.0, 90.0 | 90.0, 90.0, 90.0 | 90.0, 90.0, 120.0 |
| Resolution (Å) | 43.2 - 2.65 (2.72-2.65)[1] | 43.2 - 2.45 (2.51-2.45)[1] | 44.7 - 2.50 (2.57-2.50)[1] |
| Total Observations | 135,662 | 191,404 | 139,040 |
| Unique reflections | 20,350 | 28,936 | 13,974 |
| Redundancy | 6.7 (6.7)[1] | 6.6 (6.7)[1] | 9.9 (9.8)[1] |
| Completeness (%) | 99.8 (100.0)[1] | 99.9 (100.0)[1] | 100.0 (100.0)[1] |
| $R_{meas}$ | 0.10 (2.11)[1] | 0.07 (1.85)[1] | 0.07 (2.18)[1] |
| $I/\sigma(I)$ | 10.9 (0.9)[1] | 14.0 (1.2)[1] | 18.6 (1.0)[1] |
| $CC_{1/2}$ | 1.00 (0.57)[1] | 1.00 (0.49)[1] | 1.00 (0.48)[1] |
| **Refinement** | | | |
| Resolution (Å) | 43.2 - 2.65 | 43.2 - 2.45 | 44.7 - 2.50 |
| Reflections (work) | 16,916 | 25,465 | 12,425 |
| Reflections (test) | 1,167 | 1,466 | 659 |
| $R_{work}$ / $R_{free}$ (%) | 26.0 / 29.9 | 23.5 / 26.7 | 19.3 / 21.9 |
| Average B-factor (Å$^2$) | 113.7 | 91.4 | 89.7 |
| No. atoms | | | |
| Protein | 4,492 | 4,565 | 1,399 |
| Peptide | 88 | 0 | 0 |
| Water | 0 | 0 | 0 |
| Ramachandran[2] | | | |

| | | | |
|---|---|---|---|
| Favored (%) | 98.9 | 99.1 | 98.2 |
| Outliers (%) | 0.0 | 0.0 | 0.0 |
| Rotamers[2] | | | |
| Favored (%) | 99.2 | 97.6 | 97.2 |
| Outliers (%) | 0.4 | 0.9 | 0.7 |
| R.m.s. deviations | | | |
| Bond lengths (Å) | 0.001 | 0.002 | 0.012 |
| Bond angles (°) | 0.294 | 0.339 | 1.353 |
| Molprobity[2] | | | |
| Molprobity Score | 1.04 | 0.94 | 1.28 |
| Percentile | 100th | 100th | 100th |
| Clashscore | 2.58 | 1.84 | 5.24 |
| Percentile | 100th | 100th | 99th |
| PDB ID | 7UDM | 7UDN | 7UDO |

[1] Numbers in parentheses refer to the highest resolution shell

[2] As reported by MolProbity[4].

**Crystallization and structure determination for RPB_PEW3_R4-PAWx4**

Purified RPB_PEW3_R4 protein + PAWx4 peptide at a concentration of 36 mg/mL was used to conduct sitting drop, vapor-diffusion crystallization trials using the JCSG Core I-IV screens (NeXtal Biotechnologies). Crystals of RPB_PEW3_R4-PAWx4 grew from drops consisting of 100 nL protein plus 100 nL of a reservoir solution consisting of 0.1 M MES pH 5.0, 30% (w/v) PEG 6K at 4°C, and were cryoprotected by supplementing the reservoir solution with 5% ethylene glycol. Native diffraction data was collected at APS beamline 23-ID-D, indexed to $P2_12_12_1$ and reduced using XDS[5] (Table S1). The structure was phased by molecular replacement using Phaser[6]. A set of ~50 lowest energy predicted models from Rosetta were used as search models. Several of these models gave clear solutions, which were adjusted in Coot[7] and refined using Phenix[8]. Model refinement in $P2_12_12_1$ initially resulted in unacceptably high values for $R_{free}$ - $R_{work}$. Refinement was therefore first performed in lower symmetry space groups (P1 and $P2_1$). In the late stages of refinement, these P1 and $P2_1$ models were refined against the $P2_12_12_1$, which ultimately yielded acceptable, albeit somewhat higher R-factors.

**Crystallization and structure determination for RPB_PLP3_R6-PLPx6**

Purified RPB_PLP3_R6 protein + PLPx4 peptide at a concentration of 70 mg/mL was used to conduct sitting drop, vapor-diffusion crystallization trials using the JCSG Core I-IV screens (NeXtal Biotechnologies). Crystals of RPB_PLP3_R6-PLPx6 grew from drops consisting of 100 nL protein plus 100 nL of a reservoir solution consisting of 2.4 M (NH4)2SO4, 0.1 M Na3Cit pH4 at 18 °C, and were cryoprotected by supplementing the reservoir solution with 2.2M sodium malonate pH4. Native diffraction data was collected at APS beamline 23-ID-D, indexed to I422 and reduced using XDS[5] (Table S1). The structure was phased by molecular replacement using Phaser[6]. A set of ~28 lowest energy predicted models from Rosetta were used as search models. Several of these models gave clear solutions, which were adjusted in Coot[7] and refined using Phenix[8].

**Crystallization and structure determination for RPB_LRP2_R4-LRPx4**

Purified RPB_LRP2_R4 protein + LRPx4 peptide at a concentration of 21.4 mg/mL was used to conduct sitting drop, vapor-diffusion crystallization trials using the JCSG Core I-IV screens (NeXtal Biotechnologies). Crystals of RPB_LRP2_R4-LRPx4 grew from drops consisting of 100 nL protein plus 100 nL of a reservoir solution consisting of 0.1 M HEPES pH7, 10 %(w/v) PEG6000 at 18 °C, and were cryoprotected by supplementing the reservoir solution with 25 % Ethylene glycol. Native diffraction data was collected at APS beamline 23-ID-B, indexed to P32 2 1 and reduced using XDS[5] (Table S1). The structure was phased by molecular replacement using Phaser[6]. The coordinates of apo RPB_LRP2_R4 from the proteolyzed/filament structure were used as a search model. The resulting model was adjusted in Coot[7] and refined using Phenix[8]. Like the apo structure, this crystal structure of RPB_LRP2_R4 also contained “infinitely long filaments in the crystal, this time with peptide bound.

**Crystallization and structure determination for RPB_PLP1_R6-PLPx6**

Purified RPB_PLP1_R6 protein + PLPx6 peptide at a concentration of 143 mg/mL was used to conduct sitting drop, vapor-diffusion crystallization trials using the JCSG Core I-IV screens (NeXtal Biotechnologies). Crystals of RPB_PLP1_R6-PLPx6 grew from drops consisting of 100 nL protein plus 100 nL of a reservoir solution consisting of 0.2 M NaCl, 20 %(w/v) PEG3350 at 4 °C, and were cryoprotected by supplementing the reservoir solution with 15 % Ethylene glycol. Native diffraction data was collected at APS beamline 23-ID-B, indexed to H32 and reduced using XDS[5] (Table S1). The structure was phased by molecular replacement using Phaser[6]. A set of ~230 lowest energy predicted models from Rosetta were used as search models. Several of these models gave clear solutions, which were adjusted in Coot[7] and refined using Phenix[8]. In the later stages of refinement, two copies of the 6xPLP peptide were built into clearly defined electron density in the asymmetric unit. The first copy adopts the expected location based on the design, and makes the designed interactions with RPB_PLP1_R6. The density for this peptide and the final atomic model (19 amino acid residues) are slightly longer than the peptide used in crystallization (18 residues); this is likely due to “slippage”/misregistration of the peptide relative to the R6PO11 in many unit cells, resulting in density longer than the peptide itself. A second copy of the peptide lies across a 2-fold symmetry axis at ~50% occupancy, resulting in the superposition of this peptide with a symmetry-derived copy of itself running in the opposite direction. Despite this, the locations of each Pro/Leu side chain unit was reasonably well defined. However, it seems unlikely that the binding of the peptide at this second site would occur readily in solution.

**Crystallization and structure determination for RPB_PLP1_R6, alternative conformation 1**

Purified RPB_PLP1_R6 protein + PLPx6 peptide at a concentration of 166 mg/mL was used to conduct sitting drop, vapor-diffusion crystallization trials using the JCSG Core I-IV screens (NeXtal Biotechnologies). Crystals of RPB_PLP1_R6-PLPx6 grew from drops consisting of 100 nL protein plus 100 nL of a reservoir solution consisting of 0.02 M CaCl2, 30 % (v/v) MPD, 0.1 M NaAcet pH 4.6 at 18 °C, and were cryoprotected by supplementing the reservoir solution with 5 % MPD. Native diffraction data was collected at APS beamline 23-ID-B, indexed to P22121 and reduced using XDS[5] (Table S1). The structure was phased by molecular replacement using Phaser[6], using the coordinates for R6PO11 (alternative conformation 1) as a search model. The model was adjusted in Coot[7] and refined using Phenix[8]. In the later stages of refinement, one copy of the 6xPLP peptide was model at a site of crystal contact, where it is sandwiched between adjacent subunits in a way that is likely only bound in the crystal lattice

**Crystallization and structure determination for RPB_PLP1_R6, alternative conformation 2**

Purified RPB_PLP1_R6 protein +PLPx6 peptide at a concentration of 166 mg/mL was used to conduct sitting drop, vapor-diffusion crystallization trials using the JCSG Core I-IV screens (NeXtal Biotechnologies). Crystals of RPB_PLP1_R6-PLPx6 grew from drops consisting of 100 nL protein plus 100 nL of a reservoir solution consisting of 40 % (v/v) MPD, 0.1 M Na Phos Cit pH 4.2 at 18 °C, and were cryoprotected by supplementing the reservoir solution. Native diffraction data was collected at APS beamline 23-ID-B, indexed to P22121 and reduced using XDS[5] (Table S1). Initial attempts to phase by molecular replacement using Phaser[6] and ~500 predicted models from Rosetta and RoseTTAfold failed to yield any clear solutions. Similarly, several thousand truncations of these models (containing all combinations of 1, 2, 3, 4, or 5 of the 6 repeat units), also failed to give clear solutions. To try to identify correct but low scoring solutions in the output of these trials, we ran SHELXE autobuilding and density modification on a large number of these potential solutions. Ultimately, we were able to identify an MR solution with 2 of 6 repeats correctly placed that allowed the autobuilding of a polyalanine model and an interpretable map, which could be further improved by iterative rounds of rebuilding in Coot[7] and refinement using Phenix[8]. Ultimately, the final model revealed that in this crystal form and a similar crystallization condition (RPB_PLP1_R6, alternative conformation 1, above), RPB_PLP1_R6 adopted an alternative fold.

**Crystallization and structure determination for RPB_LRP2_R4**

Purified RPB_LRP2_R4-LRPx4 protein at a concentration of 33 mg/mL was used to conduct sitting drop, vapor-diffusion crystallization trials using the JCSG Core I-IV screens (NeXtal Biotechnologies). Crystals of RPB_LRP2_R4 grew from drops consisting of 100 nL protein plus 100 nL of a reservoir solution consisting of 0.2 M K2HPO4, 20% (w/v) PEG 3350 at 18°C, and were cryoprotected by supplementing the reservoir solution with 15% ethylene glycol. Native diffraction data was collected at APS beamline 23-ID-B, indexed to P 32 2 1 and reduced using XDS[5] (Table S1). The structure was phased by molecular replacement using Phaser[6]. A set of ~50 lowest energy predicted models from Rosetta, as well as a variety of truncated models, were used as search models. Several of these models gave clear solutions, which were adjusted in Coot[7] and refined using Phenix[8]. Four helical repeat modules were present in the asymmetric unit. However, unexpectedly, side chain density for all four repeats were very similar to one another and matched the sequence of the internal helical repeats, but not the N- and C-terminal capping repeats, which are slightly different from the internal ones. In addition, these four repeat units pack tightly against adjacent, symmetry-related molecules such that they form an “infinitely long” repeat protein running throughout the crystal. Careful examination of the the junction between each repeat unit revealed no clear breaks in electron density; the density for the backbone is continuous through the asymmetric unit, and continuous with the symmetry related molecules near the N- and C-terminus of the molecule in the asymmetric unit. Rather than truly forming an infinitely long polymer, we suspect that proteolytic cleavage of the RPB_LRP2_R4 (either during purification or crystallization led to the removal of the N- and/or C-terminal caps in many molecules, which could allow the internal repeats from separate molecules to polymerize to form fibers in the crystal. Heterogeneity in these cleavage products and how they assemble into the crystal lattice (misregistration) could consequently explain the “continuous” filaments of this repeat protein that we observe in these crystals.

### Cell Studies

#### Plasmids

For expression in cells, constructs were synthesized by Genescript and cloned into a modified pUC57 plasmid (Genscript) allowing mammalian expression under a EF1a promoter. Target peptides were cloned as C-terminal fusions with a linker (GAGAGAGRP) followed by EGFP. Binders were expressed as fusions with the first 34 residues of the Mas70p protein (Mito-tag), shown to efficiently relocalise proteins to mitochondria in mammalian cells[9] in N-terminal and with mScarlet in C-terminal[10]. Plasmids encoding the GFP-tagged peptide and the mscarlet-tagged binder were then cotransfected into cells.

Alternatively, for in vivo demonstration of the multiplexed binding between different peptides and their cognate binders (Fig.3F,G), bicistronic plasmids were generated expressing the binder flanked with a Mito-tag (respectively, PEX-tag, the first 66 residues of human PEX3, targeting to peroxisomes[11]) followed by a stop codon, then an Internal ribosome entry site (IRES) sequence and the target peptide tagged with EGFP (respectively mScarlet). Cells were then cotransfected with both bicistronic plasmids to express all four proteins.

#### Cells

U2OS cells (ATCC, HTB-96) were cultured in DMEM (Corning) supplemented with 10% fetal bovine serum (Gibco) and 1% Pen/Strep (Gibco) at 37ºC with 5% $CO_2$. Cells were transfected with Lipofectamine 3000 (Invitrogen) according to the manufacturer's instructions and imaged after 1 day of expression.

### Live-cell imaging

For live cell imaging (Fig.3), U2OS cells were plated on glass-bottom dishes (World Precision Instruments, FD35) coated with fibronectin (Sigma, F1141, 50 µg/ml in PBS), for 1 hours at 37°C DMEM-10% serum. Medium was then changed to Leibovitz's L-15 medium (Gibco) supplemented with 20 mM HEPES (Gibco) for live cell imaging. Imaging was performed onto a custom spinning disk confocal instrument composed of Nikon Ti stand equipped with perfect focus system, a fast Z piezo stage (ASI) and a PLAN Apo Lambda 1.45 NA 100X objective, and a spinning disk head (Yokogawa CSUX1). Images were recorded with a Photometrics Prime 95B back-illuminated sCMOS camera run in pseudo global shutter mode and synchronized with the spinning disk wheel. Excitation was provided by 488, 561 lasers (all Coherent OBIS mounted in a Cairn laser launch) and imaged using dedicated single bandpass filters for each channel mounted on a Cairn Optospin wheel (Chroma 525/50 for GFP and Chroma 595/50 for mScarlet). To enable fast 4D acquisitions, an FPGA module (National Instrument sbRIO-9637 running custom code) was used for hardware-based synchronization of the instrument, in particular to ensure that the piezo z stage moved only during the readout period of the sCMOS camera. Temperature was kept at 37ºC using a temperature control chamber (MicroscopeHeaters.Com, Brighton UK). System was operated by Metamorph.

### Immunofluorescence

For immunofluorescence of mitochondria (Fig. S7), U2OS FlpIn Trex cells were spread on glass-bottom dishes coated with fibronectin as above. Cells were washed with PBS then fixed in 4% PFA for 20 minutes at room temperature. Following fixation, cells were washed in with PBS and then permeabilized with 0.1% Triton X-100 in PBS for 5 minutes at room temperature. Cells were washed again with PBS and blocked in 1% BSA in PBS for 15 minutes. Cells were then incubated with TOM20 antibody (santa cruz), diluted in 1% BSA in PBS, for 1 hour at room temperature. Cells were washed 3 times with PBS and then incubated with DAPI and anti-mouse Alexa Fluor 488, diluted at 1/500e in 1% BSA in PBS, for one hour at room temperature. Cells were washed a final 3 times in PBS and then imaged using the spinning disk confocal described above.

Pull down of endogenous proteins from extracts using designed binders

For pull down of endogenous ZFC3H1 from human cell extracts, HeLa FlpIn Trex cells were lysed in lysis buffer (25 mM HEPES, 150 mM NaCl, 0.5% Tx100, 0.5% NP-40, 20 mM imidazole, pH 7.4 supplemented with Roche EDTA free protease inhibitor tablets). Lysate was incubated on ice for 10 minutes to continue lysis and then were spun at 4000 x *g* for 15 minutes at 4°C. The supernatant was incubated with pre-washed Ni-NTA agarose (Qiagen, 30210 318/AV/01) for 1 hour with rocking at 4°C to remove/reduce proteins in the lysate that bind to the resin non-specifically. For each condition, 50 µl of fresh Ni-NTA agarose resin was washed twice in lysis buffer. Equimolar amounts of purified his-tagged binder, or as a control an equal volume of buffer, was added to the Ni-NTA agarose. The pre-cleared HeLa lysate was split evenly between the 3 conditions. An input was taken of each condition, and the tubes were incubated for 2 hours at 4°C with rocking. Beads were then washed twice in lysis buffer and twice in wash buffer (25 mM HEPES, 150 mM NaCl, 20 mM imidazole pH 7.4). Proteins were then eluted from the beads in elution buffer (25 mM HEPES, 150 mM NaCl, 500 mM imidazole, pH 7.4). Inputs and elutions were ran on a NuPage 3-8% Tris-Acetate gel (Invitrogen, EA0375) and transferred to a nitrocellulose membrane using the iBlot system (ThermoFischer). Membranes were blocked in 5% (w/v) milk in TBS-TWEEN (10 mM Tris-HCl, 120 mM NaCl, 1% (w/v) TWEEN20, pH 7.4) for 30 mins at room temperature with gentle shaking. Rabbit anti-ZFC3H1 (Sigma, HPA007151, used at 1:250) and Mouse anti-alpha tubulin 488 (Clone DMA1, Sigma T6199, directly labelled with Abberior® STAR 488, NHS ester leading to a 4.5 dye/antibody degree of labelling, and used at 0.1 µg/mL final concentration) were diluted 1% (w/v) milk in TBS-TWEEN and incubated with the membrane overnight at 4°C with gentle shaking. The membrane was washed 3x in TBS-TWEEN then incubated with goat anti-Rabbit Alexa 555 (Invitrogen, A32732, 1:2000) for one hour at room temperature with gentle shaking. The membrane was washed twice with TBS-TWEEN and a final wash with TBS-TWEEN with 0.001% SDS. Membranes were imaged using a ChemiDoc system (BioRad). Alternatively, the same samples were analyzed using 4-12% Bis-Tris gels (Invitrogen NP0323BOX) and stained with Instant blue coomassie stain (Sigma ISB1L). Note that αZFC-high is also able to pull down endogenous ZFC3H1 from human cell extracts when 50 mM rather than 150 mM NaCl was used in all buffers (Figure S16C).

Mass Spectrometry

Each line of the polyacrylamide gel presented in Fig.6C was cut into six pieces (1-2 mm) and prepared for mass spectrometric analysis by manual *in situ* enzymatic digestion (the gel area containing the binder was ommited from the analysis to avoid saturation of the detector by overabundance of binder peptides). Briefly, the excised protein gel pieces were placed in a well of a 96-well microtitre plate and destained with 50% v/v acetonitrile and 50 mM ammonium bicarbonate, reduced with 10 mM DTT, and alkylated with 55 mM iodoacetamide. After alkylation, proteins were digested with 6 ng/μL Trypsin (Promega, UK), 0.1% Protease Max (Promega, UK) overnight at 37 °C. The resulting gel pieces were extracted with ammonium bicarbonate (100 μL, 100 mM) and ammonium bicarbonate/acetonitrile (50/50, 100 μL) before being dried down *via* vacuum. Clean-up of peptide digests was carried out with HyperSep SpinTip P-20 (ThermoScientific, USA) C18 columns, using 80% acetonitrile as the elution solvent before being dried down again. The resulting peptides were and were extracted in 0.1% v/v trifluoroacetic acid acid, 2% v/v acetonitrile. The digest was analysed by nano-scale capillary LC-MS/MS using an Ultimate U3000 HPLC (ThermoScientific Dionex, San Jose, USA) to deliver a flow of 250 nL/min. Peptides were trapped on a C18 Acclaim PepMap100 5 μm, 100 μm x 20 mm nanoViper (ThermoScientific, USA) before separation on PepMap RSLC C18, 2 μm, 100 A, 75 μm x 75 cm EasySpray column (ThermoScientific, USA). Peptides were eluted on a 90 minute gradient with acetonitrile and interfaced *via* an EasySpray ionisation source to a quadrupole Orbitrap mass spectrometer (Q-Exactive HFX, ThermoScientific, USA). MS data were acquired in data dependent mode with a Top-25 method, high resolution scans full mass scans were carried out (R = 120,000, *m/z* 350 – 1750) followed by higher energy collision dissociation (HCD) with collision energy 27 % normalised collision energy. The corresponding tandem mass spectra were recorded (R=30,000, isolation window *m/z* 1.6, dynamic exclusion 50 s). LC-MS/MS data were then searched against the Uniprot human proteome database, using the Mascot search engine programme (Matrix Science, UK)[18]. Database search parameters were set with a precursor tolerance of 10 ppm and a fragment ion mass tolerance of 0.1 Da. One missed enzyme cleavages were allowed and variable modifications for oxidation, carboxymethylation, and phosphorylation. MS/MS data were validated using the Scaffold programme (Proteome Software Inc., USA)[19]. All data were additionally interrogated manually. To generate the Venn diagram in Fig.6F, we considered a threshold of minimum 5 peptides to consider that a protein had been identified.

### Design Methods

### DHR Scaffolds generation

Each Designed Helical Repeat (DHR) scaffold is formed by a helix-loop-helix-loop topology that is repeated four or more times[12]. The helices range from 18 to 30 residues and the loops from 3 to 4 residues. The DHR design process goes through backbone design, sequence design and computational validation by energy landscape exploration. To match the peptides the designs were required to have a twist (omega) between 0.6 and 1.0 radians, a radius of 0 to 13 Å and a rise between 0 and 10 Å. The geometry of a repeat protein can be described by the radius of the super-helix, the axial displacement and the twist[13].

The backbone is designed using Rosetta fragment assembly guided by motifs[14]. Backbone coordinates are built up through 3,200 Monte Carlo fragment assembly steps with fragments harvested from a non-redundant set of structures from the PDB. Following each fragment insertion, the rigid body transform is propagated to the downstream repeats. The score that guides fragment assembly is composed of Van der Waal interactions, packing, backbone dihedral angles, and Residue-Pair-Transform(RPX) motifs[14]. RPX motifs are a fast way to measure the full-atom hydrophobic packability of the backbone prior to assigning side chains. After design, backbones are screened for native-like features. The loops are required to be within 0.4Å of a naturally occurring loop or rebuilt. Structures with helices above 0.14Å appear bent and kinked and are discarded. And poorly packed structures where <4 helices are in contact with each-other are filtered.

Sequence is designed using Rosetta for each backbone that passes filtering. Design begins in a symmetric mode where each repeat is identical using the RepeatProteinRelax mover. Core residues are restricted to be hydrophobic and surface residues hydrophilic using the layer design task operators. Sequence is biased toward natural proteins with similar local structure using the structure profile mover. After the symmetrical design is complete the N-terminal and C-terminal repeats are redesigned to eliminate exposed hydrophobics. Designs with poor core packing as measured by Rosetta holes < 0.5 are then filtered[15].

The designs are computationally validated using the Rosetta *ab initio* structure prediction on Rosetta@Home[16]. Rosetta *ab initio* verifies that the design is lower energy state than the thousands of alternatives conformations sampled. Simulating a protein using Rosetta@Home can take several days on hunds of CPUs. To speed this up we used machine learning to filter designs that were most likely to fail[13].

**Backbone generation of curved repeat protein monomers in Poly-Proline II conformation**

A second round of designs was made to ensure the distance between helices match the 10.9Å. distance between prolines in the Poly-Proline II. To design these backbones we used AtomPair constraints between the first helix of each repeat. The atom pair constraints were set to 10.9Å with a tolerance of 0.5 Å. For these designs we found the topologies that most efficiently produced structures that matched the AtomPair constraints had a helix length of 20 or 21 residues and a loop range of three residues.

**Peptide binders design**

**Modular peptide docking and Hashing**

To construct hash tables storing the pre-computed privileged residue interactions, we first survey the non-redundant PDB database and extract the intended interacting residues as seeds. For each seeding interaction residue pairs, random perturbations were applied to search for alternative relative conformations of the interacting residues. In the case of the sidechain-backbone bidentate interactions, the backbone residues were applied a random rigid body perturbations with a random set of euler angles drawing from a normal distribution with 0° as the mean and 60° as the standard deviation, as well as a random set of translation distances in 3D space drawing from a normal distribution with 0 Å as the mean and 1 Å as the standard deviation. At the same time, the backbone torsion angles Φ and Ψ of the backbone residue were randomly modified to values draw from a ramachandran density plot based on structures from PDB database. The transformed set of residues losing the intended interactions were discarded. The transformed residues keeping the interactions will be collected. Then the side chains of the sidechain residues were replaced with all reasonable rotamers, to further diversify the samples of the sets of interacting residues. Finally, the geometry relationship of each set of residues keeping the intended interactions were subjected to an 8D hash function (6D rigid body transformation plus two torsion angles), and represented with a 64 bit unsigned integer as the key of an entry in the hash table. The identity and the side chain torsion angles (Xs) of the side chain residues were treated as the value of the entry in the hash table. The similar processes were carried out to build different hash tables for various interactions, with minor alterations. For example, for pi-pi and cation-pi interactions, only a 6D hash function was used, because there is no need for the perturbation and consideration of the backbone torsions. For ASN, GLN, ASP or GLU interacting two residues on the backbone, a 10D hash table was applied for representing the geometry relationship, and in these cases, the geometries of the N-H and C=O groups on the backbone were treated as 5D rays.

To sample repeat peptides matching the superhelical parameters of the DHRs, we randomly generate a set of backbone torsion angles φ and ψ, for example, [$\varphi_1$, $\psi_1$, $\varphi_2$, $\psi_2$, $\varphi_3$, $\psi_3$] for repeats of tri-peptide. If there any pair of φ and ψ angles get a high

Rosetta Ramachandran score above the threshold -0.5, it means that this pair of torsion angles are likely to introduce intra-peptide steric clashes, and we will randomly regenerate a new pair of φ and ψ angles until they are reasonable according to the Rosetta Ramachandran score. Next, we will set the backbone torsion angles of the repeat peptide using this set of φ and ψ angles repetitively across the 8-repeats. And we will calculate the superhelical parameters using the 3D coordinates of adjacent repeat units of the repeat peptide. The repeat peptides matching the superhelical parameters of any one of the curated DHRs, will be saved for the docking step.

To dock cognate repeat proteins and repeat peptides, with matching superhelical parameters, they are first aligned to the z axis by their own superhelical axes. Next step, a 2D grid search (rotation around and translation along the z axis) were carried out to sample compatible positions of the repeat peptide in the binding groove of the repeat protein. Once a reasonable dock is generated without steric clash, the relevant hash function will be used to iterate through all potential peptide-protein interacting residue sets, to calculate the hash keys. If a hash key exists in the hash table, the interacting side chain identities and torsion angles would be pulled out immediately and installed on all equivalent positions of this repeat peptide-repeat protein docking conformation. The docked peptide-DHR pair will be saved for the interface design step if the peptide-DHR hydrogen-bond interactions are satisfied.

**Peptide binding interface design**

If a single dock was accepted with the designed repetitive peptide-DHR hydrogen-bond, the peptide was first trimmed to the exact same repeat number as the DHR (e.g., 4-repeat or 6-repeat). After that, for both peptide and DHR sides, each amino acid was set linked to its corresponding amino acids on the same position in each repeat unit. This was to make sure all the following design steps would be carried out with the exact same symmetry inside of both the DHR and peptide.

During our design cycles, the interface neighbor distance is set as 9Å as the whole designable range around the DHR-peptide binding interface, and 11Å as the whole minimization range. Three rounds of full hydrophobic fastdesign[14] followed by hydropathic fastdesign were carried out, with each hydrophobic or hydrophilic fastdesign repeating twice. The Rosetta score function beta_nov16 was chosen in all design cycles. In the produced complex, the peptide itself with an averaged score (three calculations were carried out) larger than 20.0 or the complex scored larger than -10.0 were rejected directly.

After the preliminary design was done, we carried out two types of sanity checks to further optimize the designed peptide sequence, as well as the designed DHR interface. Specifically, for the peptide side, in the tri-peptide repeat units, every two amino acids other than Proline were scanned for a possible mutation to all twenty amino acids except cysteine, unless a certain originally designed peptide amino acid is making the hashed sidechain-backbone hydrogen-bond, or sidechain-sidechain hydrogen-bond, or sidechain-sidechain-backbone hydrogen-bond with the DHR interface. DDG (binding energy for the peptide-DHR complex) was compared before and after this peptide side mutation; and the mutation was accepted if the delta DDG (DDG_after - DDG_before) was larger than 1.0. Similarly, we also checked the designed DHR interface by mutation. The whole DHR was scanned. For the designed hydrophobic amino acids which were originally hydrophilic, a delta DDG of -5.0 was set as the threshold to be accepted as a necessary design which made enough binding contribution. For the designed hydropathic amino acids, a delta DDG of -2.0 was used as threshold.

For experimental characterization, we selected designed complexes with near ideal bidentate hydrogen bonds between protein and peptide, favorable protein-peptide interaction energies (DDG <= -35.0), interface shape complementary (Iface_SCval >= 0.65), tolerable interface unsatisfied hydrogen bonds (Iface_HbondsUnsatBB <= 2, Iface_HbondsUnsatSC <=4) and low peptide apo energies (ScoreRes_chainB <= 0.9).

Forward Docking

As for the selected designed complexes from our round-two experiments, forward docking was performed to ensure the specificity in silico. For each designed complex, 10,000 arbitrary peptide conformations were generated as above, using the designed sequence. The same docking protocol was conducted as described in the docking stage, against the untouched designed DHR. Fastrelax[17] was then performed for the 10,000 docks, and the DDG vs. peptide backbone RMSD was plotted to check the convergence of the complex. Only the “converged” complexes were selected for experimental characterization, e.g., i) peptide backbone RMSD<2.0Å among the top 20 designs with lowest DDG during Forward Docking and ii) the averaged peptide backbone of the top 20 designs was close to the original design model (RMSD<1.5Å).

SSM library preparation

We carried out the site saturation mutagenesis (SSM) studies for some of the designed peptide-protein binding pairs to gain better understanding of the peptide binding modes, and to search for improved peptide binders. For each designed repeat protein, we ordered a SSM library covering the central span of 65 amino acids within the whole repeat protein, due to the chip DNA size limitation. This span roughly equals one and half repeating units, across three helices. The chip synthesized DNA oligos for the SSM library then amplified and transformed to EBY100 yeast together with a linearized pETCON3 vector including the encoding regions of the rest of the designed repeat protein. Each SSM library was subject to an expression sort first, in which the low-quality sequences due to chip synthesize defects or recombination errors were filtered out. The collected yeast population, which successfully expresses the designed repeat protein mutants, will be re-grown, and be subject to the further round of peptide binding sorts. The next-generation sequencing results of this yeast population will also serve as the reference data for SSM analysis. The next round of without-avidity peptide binding sorts used various concentrations of the target peptide, depending on the initial peptide binding abilities, ranging from 1 nM to 1000 nM. The peptide-bound yeast populations were collected and sequenced by using Illumina NextSeq kit. The mutants were identified and compared to the mutants in the expression libraries. The enrichment analysis was carried out to identify the beneficial mutants and provide information for interpreting the peptide binding modes. For each mutant, its enrichment value is calculated by dividing its ratio in peptide-bound population by its ratio in expression population. The enrichment value is then subject to a log10 transformation, and plotted in heatmaps for the SSM analysis.

### Design of binders against endogenous targets

To evaluate which endogenous proteins could be currently targeted with our method (Fig.6), we developed a python code to search databases for subsequences matching permutations of the set of amino acid triplets we designed binders for in this study (i.e LRP PEW PLP IYP PKW IRP LRT LRN LRQ RRN PSR PRQ). This code can be accessed freely (https://github.com/tjs23/prot_pep_scan). We then ranked all outputs to find the longest subsequence possible, and manually inspected the candidates to find subsequences landing in disordered regions. Doing this analysis on the human proteome suggested that ZFC3H1 could be a good target for two main reasons: 1) this protein possess the sequence (PLP)x4 within a large disordered domain, with downstream sequence (PEDPEQPPKPPF) within the reach of our binder design method and 2) the protein is well studied, and, in particular, commercial, highly specific and validated antibodies exist against it.

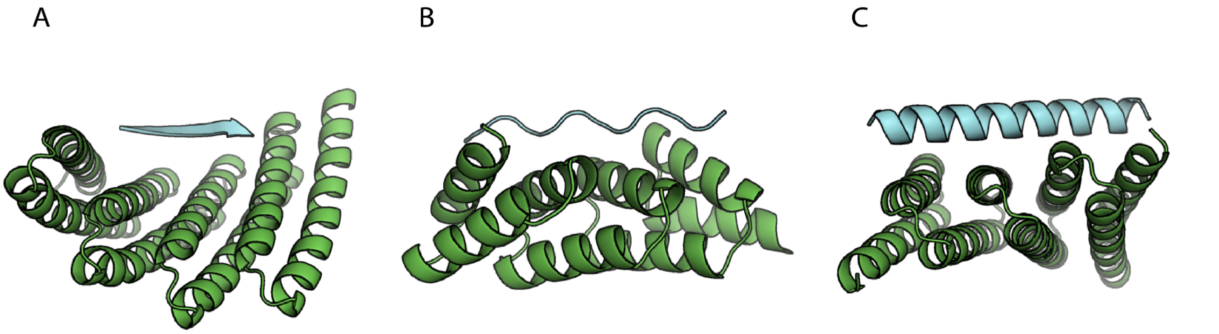


**Supplementary Figure 1:** Examples of repeat proteins computationally designed to bind to A) extended beta strand, B) polypeptide II, and C) helical peptide backbones.

| Design ID | Design target | Experimental target | di-peptide target | Tri-peptide target |
|---|---|---|---|---|
| PXX02 | PSD | PEW | | 1 |
| PXX10 | PMP | PWP/PAW | | 2 |
| PXX12 | ADP | LRP/LRT | | 2 |
| PXX27 | PDH | PDW | | 1 |
| PXX28 | PMP | PAW/PKW | | 2 |
| PXX45 | PD | DW | 1 | |
| CP06 | VRP | LRT/INR | | 2 |
| CP10 | IMP | IRP | | |
| CP25 | PSM | LRP/MYP | | 2 |
| CP33 | PKL | PRM/PRY/PKW | | 3 |
| KP02 | DI | PAN/PEW | | 2 |
| KP09 | RP | LRA | | 1 |
| KP22 | DP | PEW/EW | 1 | 1 |
| summary | | | 2 | 19 |

**Supplementary Table 2:** First-round experimental characterization summary. It is clearly shown that among the binders, most of them bound peptides with sequences similar to those targeted but not the same; and peptides with 3 residue repeat units were targeted more successfully (19 in total) than those with 2 residue repeat units (2 in total).

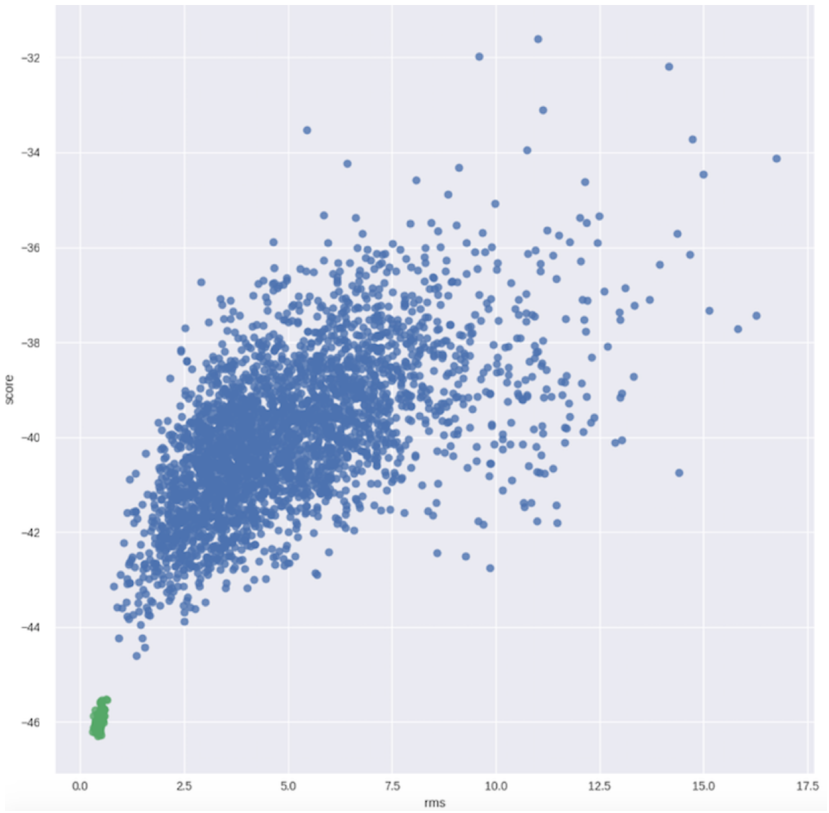


**Supplementary Figure 3:** Monte Carlo flexible backbone docking calculations after design to assess the structural specificity of the designed peptide binding interface. It started from large numbers of peptide conformations randomly generated with superhelical parameters in the range of those of the proteins (usually 10,000 - 50,000 trajectories), and selected those designs with converged peptide backbones (RMSD<2.0 among the top 20 designs with lowest DDG) close to the design model (RMSD<1.5). Green dots shown in the above example plot represent the converged designs picked by this threshold.

| Tested | Expressed well | Monomeric | Binding to the designed target by yeast surface display | Affinity < uM by yeast surface display & Octet, BLI |
|---|---|---|---|---|
| 54 | 42 | 25 | 19 | 16 |

**Supplementary Table 4:** Second-round experimental characterization summary. 54 second-round designed protein-peptide pairs were tested in total. 42 of the designed proteins were solubely expressed in E coli, 25 were monomerically dispersed by Size Exclusion Chromatography (SEC), and 16 designed bound their targets with considerably higher affinity and specificity than in the first round.

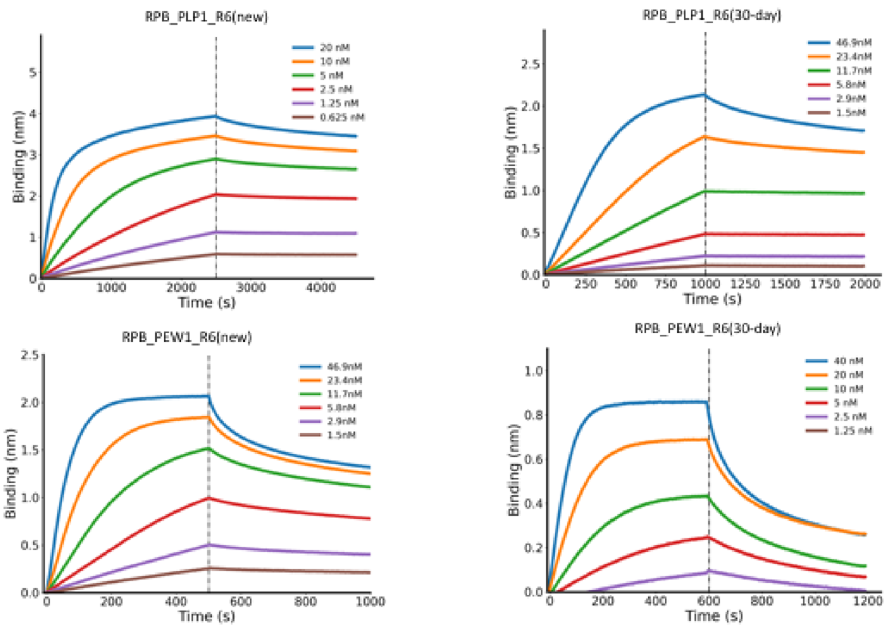


**Supplementary Figure 5:** little decrease in binding observed for designs RPB_PLP1_R6 and RPB_PEW1_R6 30-day old in 4C. Biolayer interferometry characterization of binding of designed proteins to the corresponding peptide targets. Two-fold serial dilutions were tested for each binder, and the full tested concentration is labeled. The biotinylated target peptides were loaded onto the Streptavidin (SA) biosensors, and incubated with designed binders in solution to measure association and dissociation.

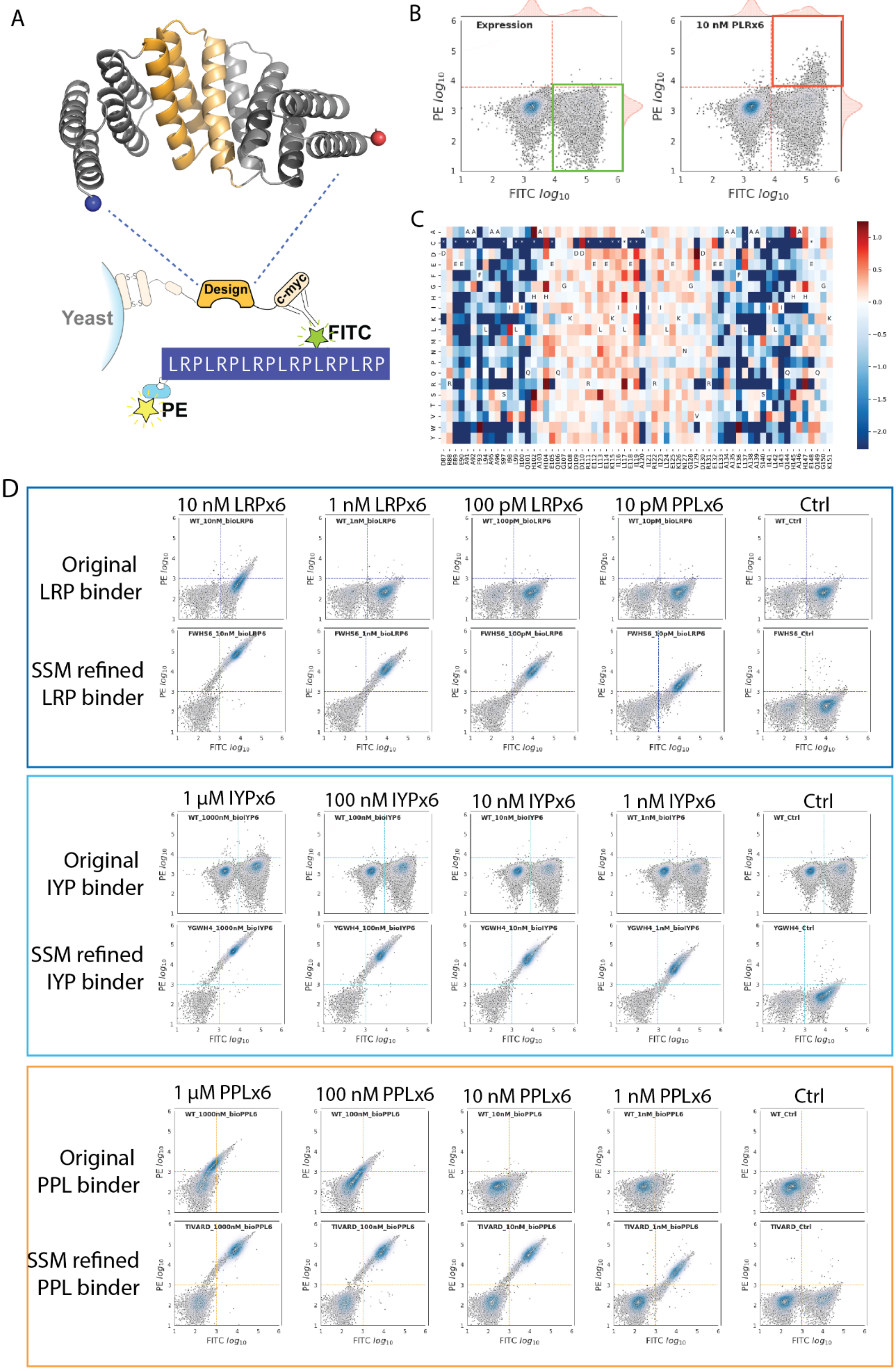
A
B
C
D
Yeast
Design
c-myc
FITC
PE
LRPLRPLRPLRPLRPLRP
Expression
10 nM PLRx6
PE $log_{10}$
FITC $log_{10}$
10 nM LRPx6
1 nM LRPx6
100 pM LRPx6
10 pM PPLx6
Ctrl
Original
LRP binder
SSM refined
LRP binder
1 µM IYPx6
100 nM IYPx6
10 nM IYPx6
1 nM IYPx6
Original
IYP binder
SSM refined
IYP binder
1 µM PPLx6
100 nM PPLx6
10 nM PPLx6
1 nM PPLx6
Original
PPL binder
SSM refined
PPL binder

**Supplementary Figure 6:** Site saturation mutagenesis libraries (SSMs) are constructed and screened for enhancing peptide binding abilities of designed repeat peptide binders. A) A schematic illustration of the mutagenesis region within the designed repeat protein, and the principles of the yeast surface display assay for peptide binding analysis. In short, the biotinylated repeat peptides (a six-repeat of LRP peptide is shown as an example) are synthesized and can be detected by SAPE, while the expression of designed protein on yeast surface are monitored by FITC conjugated anti-Myc antibody. A double high signal of both PE and FITC, using Flow Cytometry, indicates the valid peptide binding events. B) The SSM libraries are first subjected to expression sorting (Left panel), in which there is no targeted peptide added. The yeast populations, which display well expressed SSM mutants, will show above threshold FITC signals, are collected (green box) for next generation sequencing, and are re-grown for the next rounds of sorting. In the next round sorting, the targeted peptide is incubated with the yeast library, and labeled by both FITC and SAPE (Right panel). The $FITC^{+}PE^{+}$ population is collected for analysis (orange box). 3) By using next generation sequencing, enrichment analysis for each mutation is carried out, and a heatmap for all mutations is generated. In this heatmap, using a designed LRP binder SSM library as an example, the red shades indicate enrichment with incubating with the targeted peptide, and the blue shades indicate depletion. Several mutations show exceptional enhancement of the LRP repeat peptide binding ability, such as F93W, H102S, etc. 4) Using the SSM library, we can dramatically enhance the peptide binding abilities of the designed peptide binder. Three example yeast display assays titrating the peptide concentrations are shown here. The top row of each example is using the originally designed peptide binder, and the bottom row is using the peptide binder containing the combinations of the best mutations discovered in the SSM library screenings. ~1000x increase of the peptide binding ability can be achieved with the assistance of SSM libraries. Note, the ratio of yeast population in the upper right quadrant indicates the peptide binding ability.

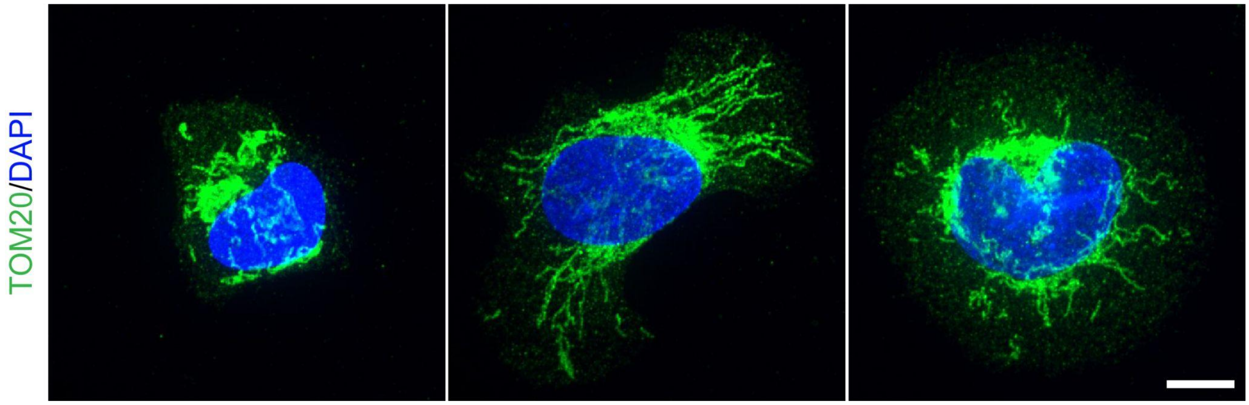


**Supplementary Figure 7: Mitochondria immunostainings in control U2OS cells**
*Wild type* U2OS cells were spread onto fibronectin coverslips as in Fig.3, then fixed and processed for immunofluorescence using TOM20 antibodies as a marker of mitochondria. Note that mitochondria appearance in these control cells is similar to that observed upon overexpression of designed binders fused to mitochondria targeting sequences (Fig.3). suggesting that these constructs do not affect mitochondria shape. Scale bar: 10 µm.

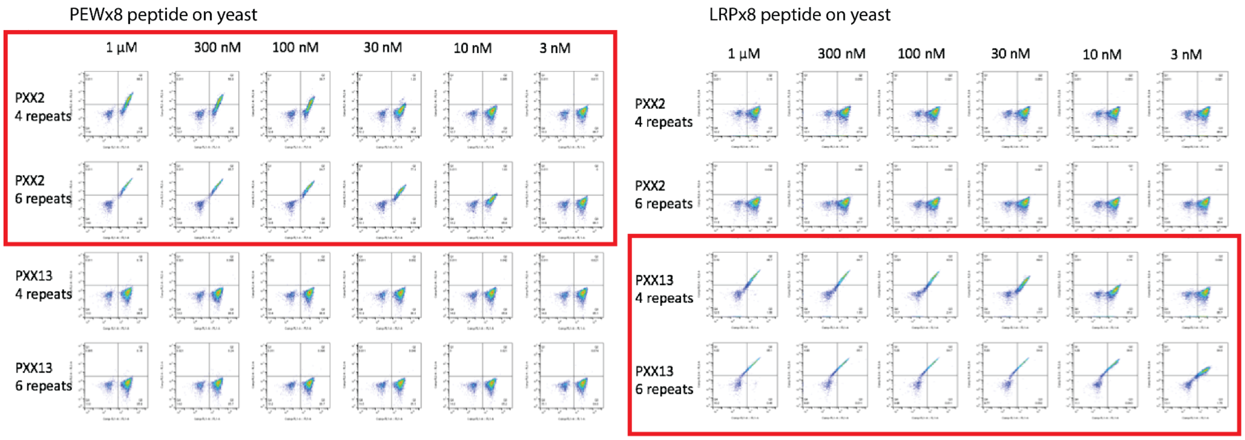


**Supplementary Figure 8:** Six-repeat versions of RPB_LRP2_R6 and RPB_PEW2_R6 had higher affinity for eight-repeat LRP and PEW peptides than four-repeat versions without any decrease in specificity. The yeast surface display assay was performed as described above. In this case, the biotinylated repeat proteins (the six-repeat of RPB_LRP2_R6, RPB_PEW2_R6 and the four-repeat version RPB_LRP2_R4, RPB_PEW2_R4) are synthesized and can be detected by SAPE, while the expression of designed repeat peptide on yeast surface are monitored by FITC conjugated anti-Myc antibody. A double high signal of both PE and FITC, using Flow Cytometry, indicates the valid peptide binding events. Serial dilutions were tested for each binder, and the full tested concentration is labeled.

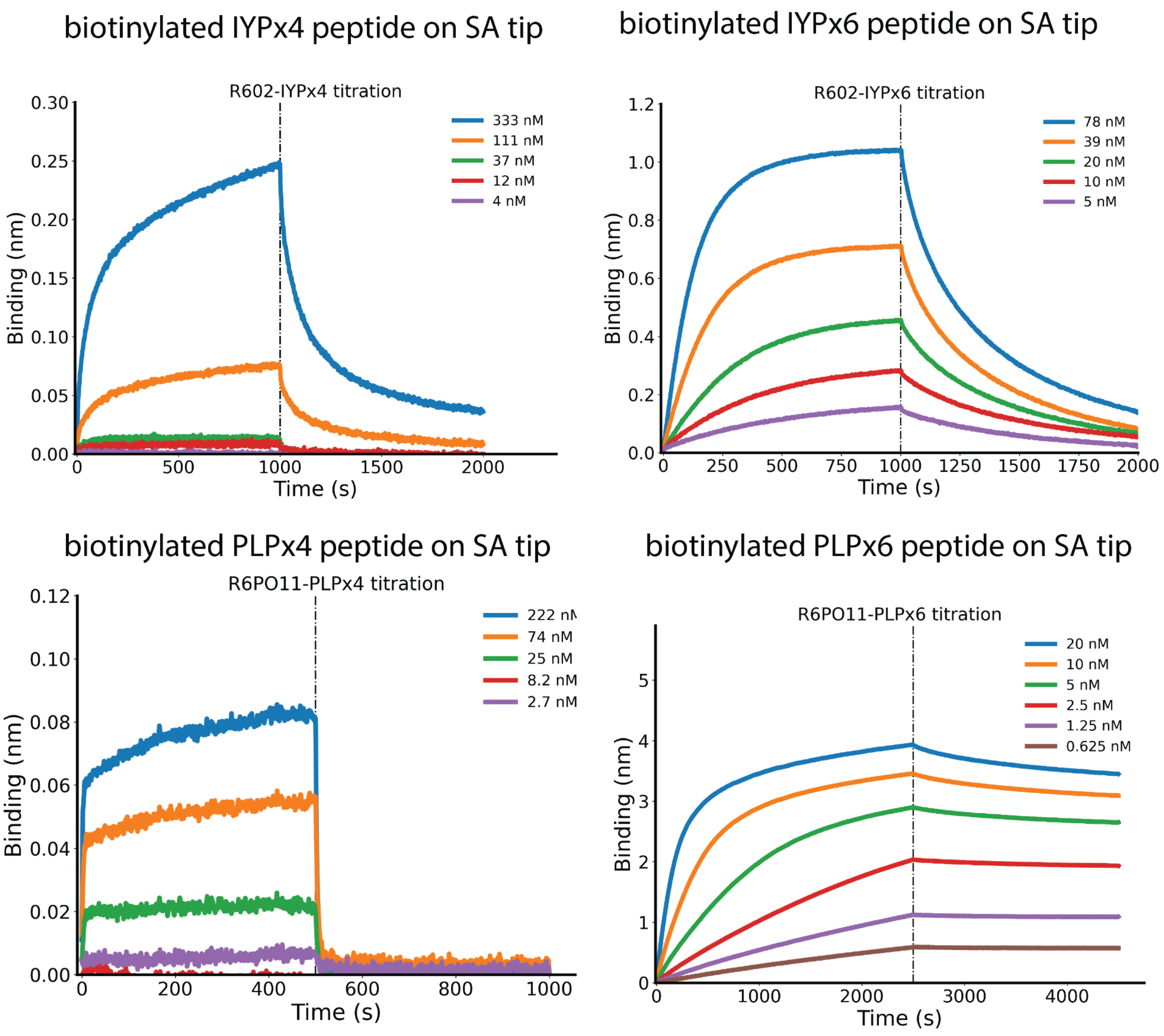


**Supplementary Figure 9:** Six-repeat IYP and PLP peptides had higher affinity for six-repeat versions of the cognate designed repeat proteins (RPB_IYP1_R6, RPB_PLP1_R6) than four-repeat versions. Biolayer interferometry characterization of binding of designed proteins to the corresponding peptide targets. Three-fold (for four-repeat peptides experiments) and two-fold (for six-repeat peptides experiments) serial dilutions were tested for each binder, and the full tested concentration is labeled. The biotinylated target peptides were loaded onto the Streptavidin (SA) biosensors, and incubated with designed binders in solution to measure association and dissociation. Particularly, the dissociation rate was obviously dramatically increased when testing against the six-repeat peptides than the four-repeat peptides, indicating a much tighter binding event.

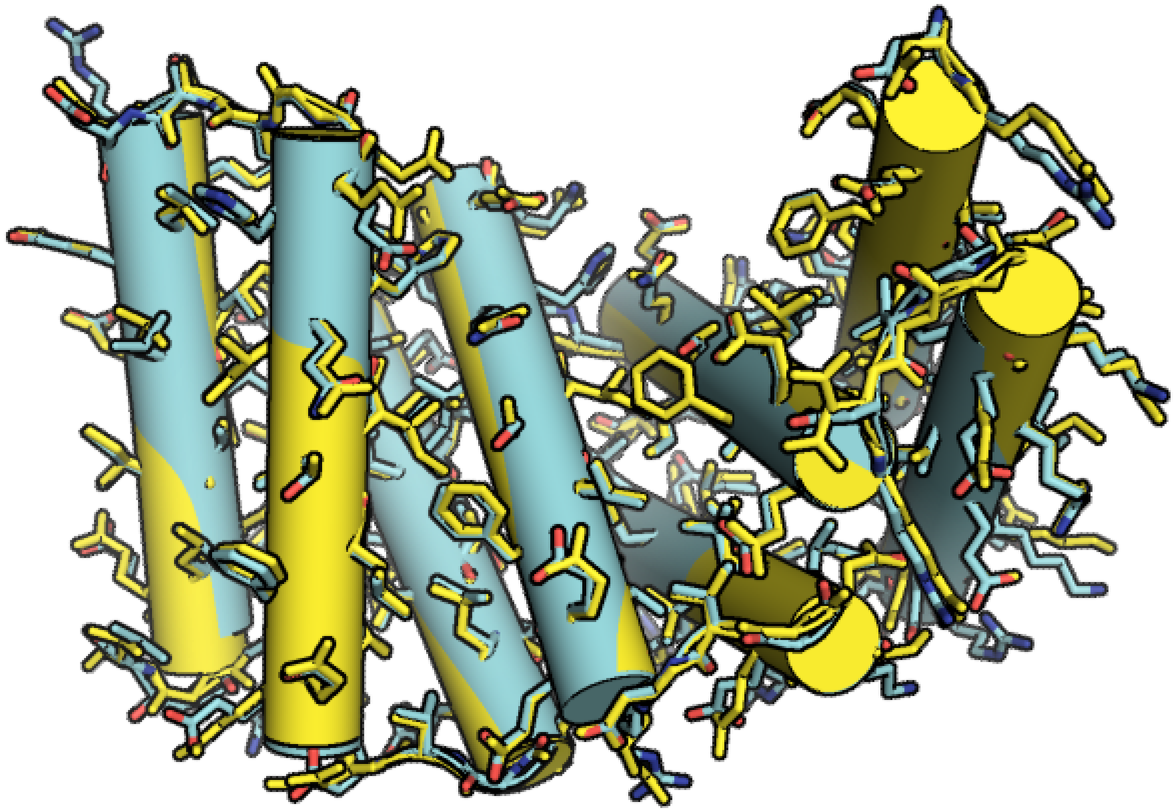

**Supplementary Figure 10:** the crystal structure of the unbound first-round design RPB_LRP2_R4 (yellow) aligned with the design model (cyan).

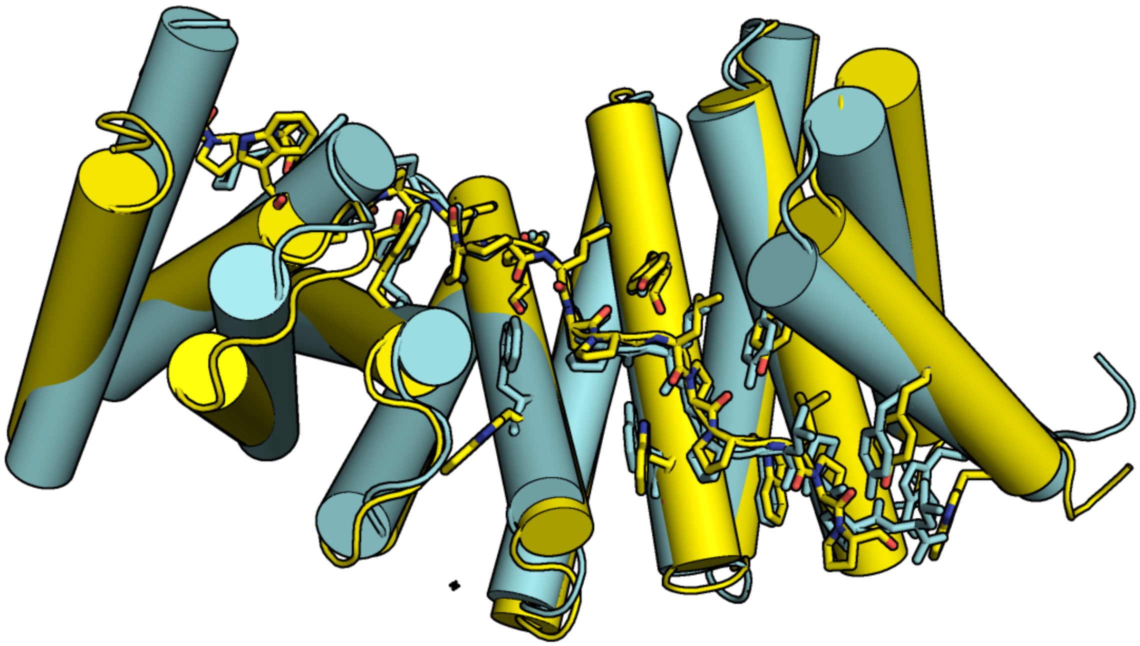

**Supplementary Figure 11:** the crystal structure of the first-round complex RPB_PLP3_R6 - PLPx6 (yellow) aligned with the design model (cyan). As is shown here, the peptide PLP units fit exactly into the designed curved groove formed by repeating tyrosine, alanine, and tryptophan residues matching the design model with near atomic accuracy, with Cα rmds of 1.70 Å for the binder apo, 2.00 Å for the peptide neighbor interface and 1.64 Å for the whole complex.

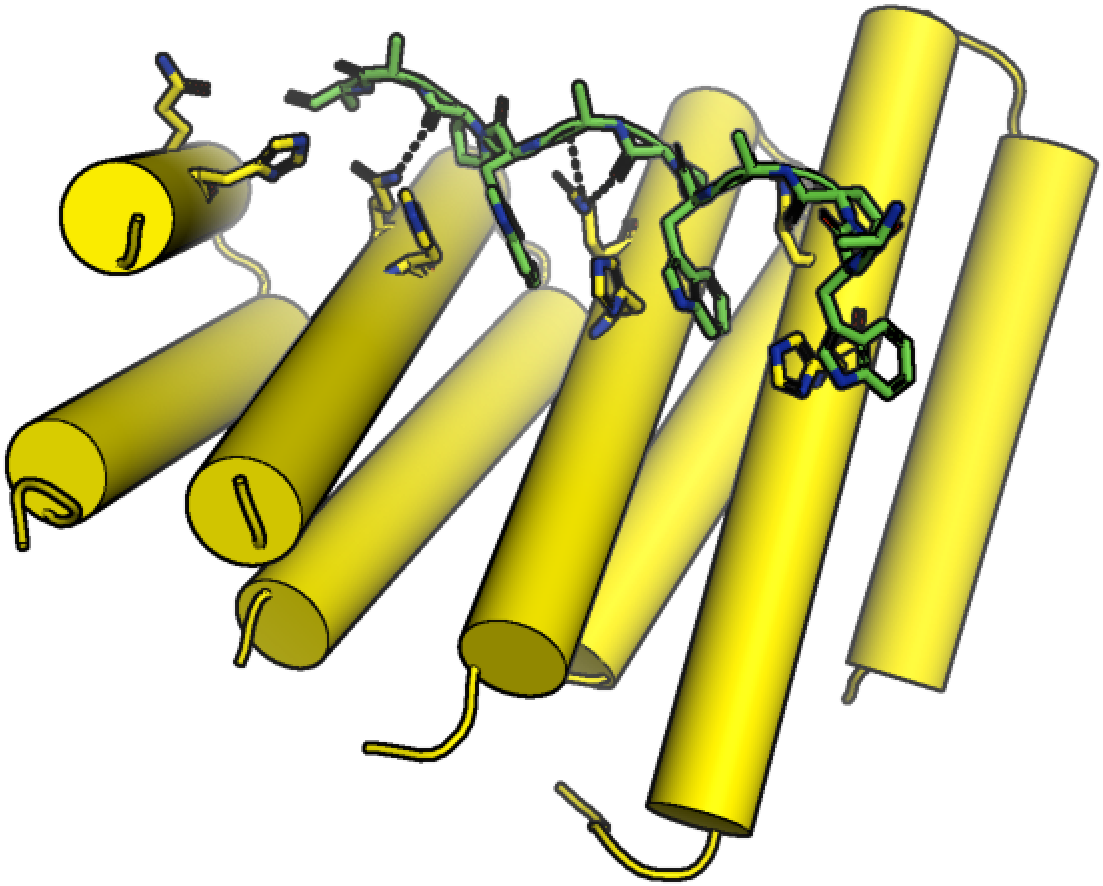

**Supplementary Figure 12:** the co-crystal structure of RPB_PEW3_R4 - PAWx4, the PAW units bind to a relatively flat groove formed by repeating histidine residues and glutamine residues as designed (shown as sticks).

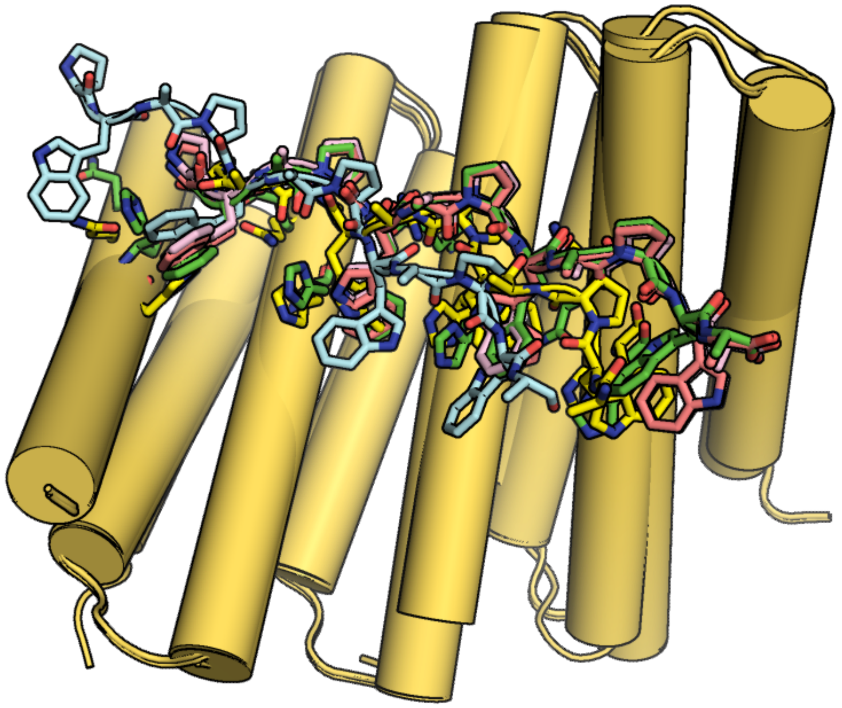

**Supplementary Figure 13:** top 5 complex PDBs from the flexible docking generated ensemble. Green, pink and gray are the ones closest to the crystal structure (shown in yellow) with RMSD over the peptide and the binding residues ~0.03 Å, while the cyan dock RMSD=3.89 Å.

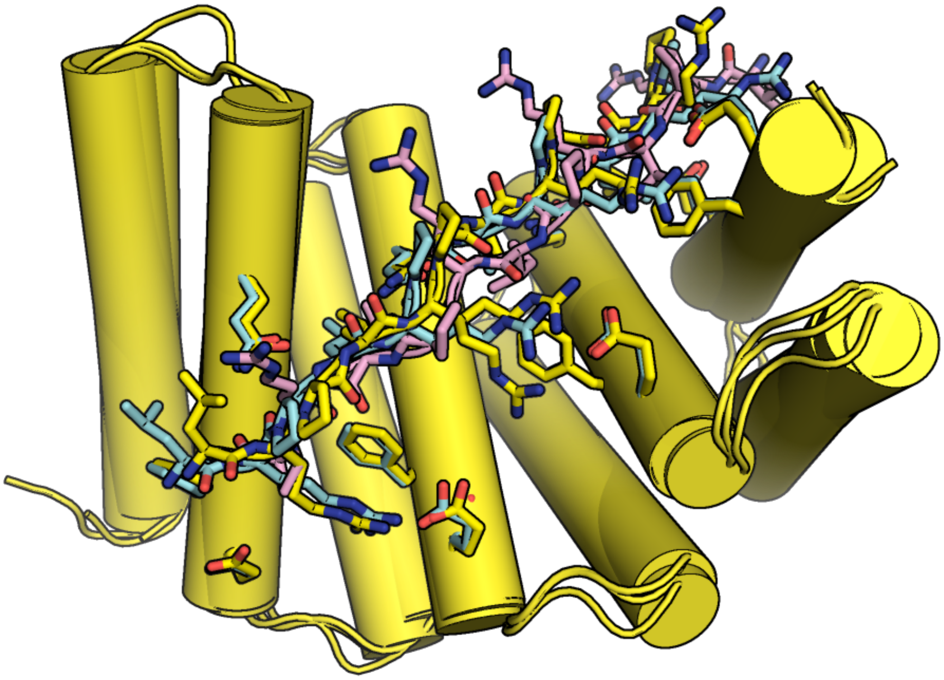

**Supplementary Figure 14:** top 5 complex PDBs from the flexible docking generated ensemble. Green, pink and gray are the ones closest to the crystal structure (shown in yellow) with RMSD over the peptide and the binding residues ~0.03 Å, while the cyan dock RMSD=3.89 Å.

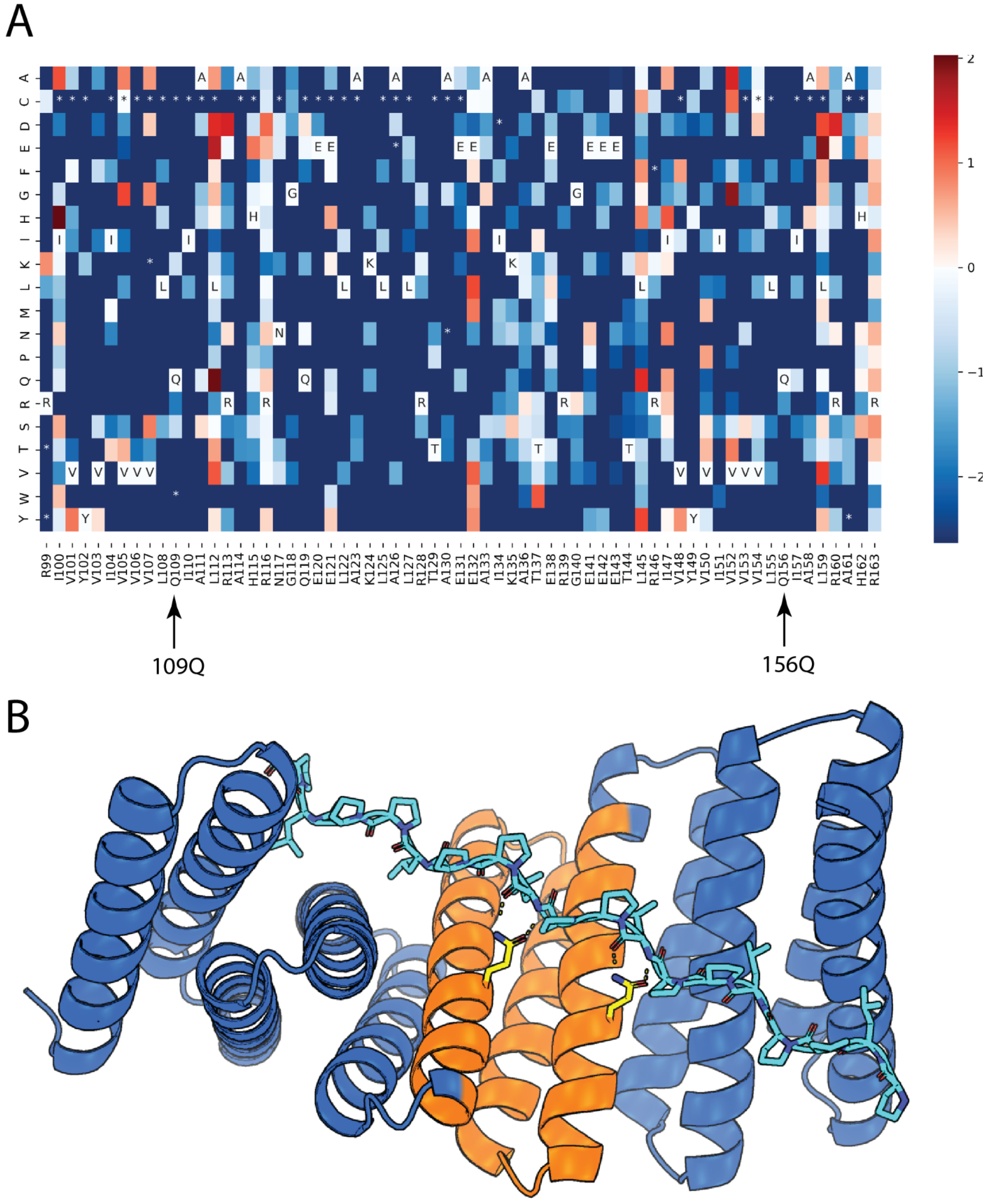


**Supplementary Figure 15:** SSM binding interface footprinting results were consistent with the design model and crystal structure. A. Using a PPL repeat peptide binder as an example, a heatmap presenting enrichment analysis for each mutation is generated. In each cell, the red color indicates enrichment, while the blue color indicates depletion. WT sequences are indicated in the cells labeled with amino acid one-letter codes. The mutants missing in the expression library are labeled with asterisks. Two positions (109Q and 156Q) are highlighted as examples showing conserved positions. Almost all mutations other than the WT in these two positions are greatly depleted. B) Illustration shows the SSM region (Orange), and the two conserved positions (109Q and 156Q in yellow).

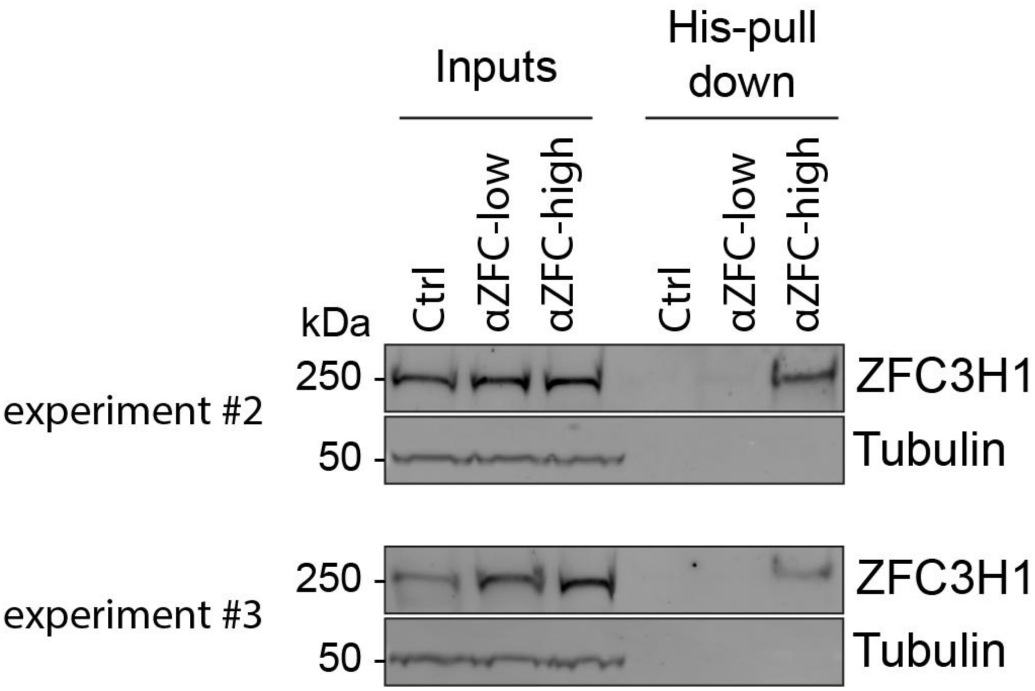


**Supplementary Figure 16: Characterization of ZFC3H1 binders**

Hela cell extracts were subjected to pull down using indicated binders bound to NiNTA agarose beads, or naked beads as control. Recovered proteins were processed for western blot against endogenous ZFC3H1 (or tubulin as a loading control). Two completely independant experiments are shown. This experiments are repeats of the experiment presented in Fig.6E, albeit at a different salt concentration, namely 50 mM instead of 150 mM.

---

| Peptide | PLPx6 | LRPx6 | PEWx6 | IYPx6 | PRMx6 | PKWx6 |
|---|---|---|---|---|---|---|

| | | | | | | |
|---|---|---|---|---|---|---|
| binder | | | | | | |
| Vr | 1.25 | 0.58 | 1.30 | 1.76 | 1.13 | 0.58 |

**Supplementary Figure 17: Structural validation of six repeat peptide binders by SAXS volatility ratio (Vr) calculation.**

| name | RPB_PLP1 _R6-PLPx6 | RPB_LRP2 _R4-LRPx4 | RPB_PLP3 _R6-PLPx6 | RPB_PEW3 _R4-PAWx4 |
|---|---|---|---|---|
| full_iRMSD | 1.61 | 1.43* | 6.49 | 12.41 |
| inter_iRMSD | 1.34 | 1.22* | 4.08 | 9.27 |

**Supplementary Figure 18: Interface side-chain heavy atom RMSD calculation for four co-crystal complexes and design models.**

The interface heavy atom RMSD calculations using pymol align with cycles=0 (iRMSD for short) was applied to all four crystal-design complexes. For the first-round designs, for example, the values are averaged over five top designs for RPB_PEW3_R4-PAWx4 since the design models were not fully converged (as stated in the main text). For RPB_LRP3_R4-LRPx4, since the final two models were sampling two distinct Arginine romaters as stated in the main text, we calculated iRMSD for these two respectively. The closest one was shown above with asterisks, and the further one as (full_iRMSD = 5.29, inter_iRMSD = 5.16). For all four pairs, we inspected both the full-repeat RMSD with internal-repeat RMSD (N-terminal and C-terminal caps excluded) here, due to the potential lever-arm effect.

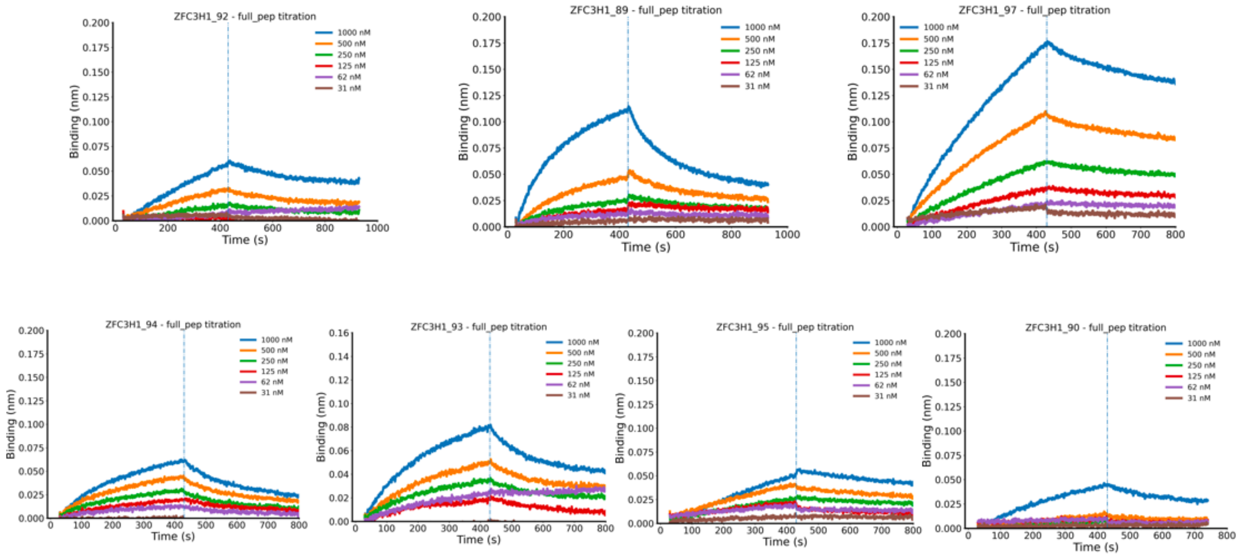


**Supplementary Figure 19: Biolayer Interferometry screening for the seven endogenous ZFC3H1 binders.**

Two-fold serial dilutions were tested for each binder, and the full tested concentration is labeled. The biotinylated target 24 amino-acid peptide (PLPPLPPLPPLPPEDPEQPPKPPF) were loaded onto the Streptavidin (SA) biosensors, and incubated with designed binders in solution to measure association and dissociation. Two tightest binders (ZFC3H1_93_b7 and ZFC3H2_97_b2) were selected for further fluorescence polarization (FP) characterization and cell assays.

**Legend to supplementary Table 5 (provided as Excel file).**
Full proteomics dataset corresponding to the data presented in Fig.6D. A minimal threshold of five peptides was considered for identification of a protein. Proteins specifically pulled down by the αZFC-high binder but absent in the beads or αZFC-low negative control binders are presented in a second table for convenience.
